# Aminobenzoic acid derivatives obstruct induced fit in the catalytic center of the ribosome

**DOI:** 10.1101/2023.01.25.525534

**Authors:** Chandrima Majumdar, Joshua A. Walker, Matthew B. Francis, Alanna Schepartz, Jamie H. D. Cate

## Abstract

The *Escherichia coli* ribosome can incorporate a variety of non-l-α-amino acid monomers into polypeptide chains, but with poor efficiency. Although these monomers span a diverse set of compounds, there exists no high-resolution structural information regarding their positioning within the catalytic center of the ribosome, the peptidyl transferase center (PTC). Thus, details regarding the mechanism of amide bond formation and the structural basis for differences and defects in incorporation efficiency remain unknown. Within a set of three aminobenzoic acid derivatives–3-aminopyridine-4-carboxylic acid (Apy), *ortho*-aminobenzoic acid (*o*ABZ), and *meta*-aminobenzoic acid (*m*ABZ)–the ribosome incorporates Apy into polypeptide chains with the highest efficiency, followed by *o*ABZ and then *m*ABZ, a trend that does not track with the nucleophilicity of the reactive amines. Here, we report high resolution cryo-EM structures of the ribosome with these three aminobenzoic acid derivatives charged on tRNA bound in the aminoacyl-tRNA site (A site). These structures reveal how the aromatic ring of each monomer sterically blocks positioning of nucleotide U2506, thereby preventing rearrangement of nucleotide U2585 and the resulting induced fit in the PTC required for efficient amide bond formation. They also reveal disruptions to the “proton wire” responsible for facilitating formation and breakdown of the tetrahedral intermediate. Together, the cryo-EM structures reported here provide a clear rationale for differences in reactivity of aminobenzoic acid derivatives relative to l-α-amino acids and each other, and point to stereochemical constraints on the size and geometry of non-proteinogenic monomers that can be accepted efficiently by wild-type ribosomes.

## Introduction

Ribosomes synthesize natural proteins by catalyzing amide bond formation between an l-α-amino acid and a l-α-peptidyl oligomer linked as esters to the 3’ termini of two tRNAs. The reaction follows an acyl addition-elimination pathway; one tRNA (the A-site tRNA) carries the α-amine nucleophile, whereas the other (the P-site tRNA) carries the carbonyl electrophile. The reaction occurs within a region of the 23S ribosomal RNA (rRNA) called the peptidyl transferase center (PTC). The PTC is composed of roughly 180 nucleotides (nts), organized into two shells, that collaboratively position the A-site α-amine nucleophile and the P-site carbonyl electrophile in close proximity, thus functioning as an “entropic trap” to enhance the rate of bond formation by more than seven orders of magnitude.^1–3^ The outer shell of the PTC, composed of nucleotides on the apical loops of rRNA helices 80 and 92, base pair with the CCA ends of the P- and A–site tRNAs and position the appended amino acid esters within the inner shell composed of universally conserved nucleotides A2451, U2506, U2585, and A2602 (*Escherichia coli* numbering).^2^ Studies of the ribosome in multiple states of catalysis indicate that bond formation is also facilitated by induced fit, in which substrate-induced conformational changes within the PTC fine-tune the relative orientation of the nucleophile and electrophile. The presence of a correctly positioned l-α-peptidyl oligomer in the P site and an α-amino acid ester in the A site within the inner shell reorients U2506 and U2585 to protect the reaction center from the side chain of the A-site monomer and position the α-amine and carbonyl moieties in line for nucleophilic attack.^4^ Bond formation is further facilitated by three ordered water molecules, a “proton-wire”, that deprotonate the α-amine nucleophile and activate the carbonyl electrophile prior to nucleophilic attack, stabilize the tetrahedral intermediate after nucleophilic attack, and reset the PTC for another round of amide bond formation.^5^

A major advance in the cellular biosynthesis of novel protein materials was the discovery that this induced fit and proximity-guided mechanism tolerates a diverse range of canonical and non-canonical l-α-amino acid side chains within the A site, the P site, and within the exit tunnel.^6^ Recently it has been shown that a number of non-l-α-amino acid monomers are also tolerated, at least *in vitro.* Examples of non-l-α-amino acid monomers that have been introduced into growing polypeptide chains include D-α-amino acids, α-hydroxy and α-thio acids, β^2^- and β^3^-amino acids, α-aminoxy and α-hydrazino acids, and a small number of aminobenzoic acid derivatives.^7–15^ Although these molecules are accepted as ribosome substrates in highly optimized *in vitro* translation reactions, the yields of polypeptide products vary significantly as a function of location, context, the presence or absence of supplementary translation factors such as EF-P, and monomer identity. Of this set, the only monomers incorporated into proteins *in vivo* by wild type ribosomes thus far are α-hydroxy acids, and the factors limiting their incorporation efficiency are unknown.^16–19^

One class of non-l-α-amino acid monomers evaluated as substrates for wild type ribosomes includes a variety of differentially substituted aminobenzoic acid derivatives.^13^ These monomers are of great interest as both materials and pharmaceuticals. Aromatic polyamides (aramids) represent a fundamental building block of both heat- and impact-resistant fabrics as well as many microbial natural products, fold into unique secondary structures including β-turns and helices, and can improve the drug-like properties of cyclic peptides.^20–23^ Although certain aminobenzoic acid monomers can be introduced into short peptide oligomers by the ribosome, the incorporation efficiencies vary widely. Specifically, 3-aminopyridine-4-carboxylic acid (Apy) acid was incorporated into short peptide oligomers almost 5-fold more efficiently than *ortho*-aminobenzoic acid (*o*ABZ), whereas analogous oligomers containing *meta*-aminobenzoic acid (*m*ABZ) were barely detected by mass spectrometry.^13,16^

We hypothesized that the reduction in ribosomal incorporation efficiency of aminobenzoic acid derivatives compared to l-α-amino acids, and the differences in yields between the aminobenzoic acid derivatives, may be related to their positioning within the PTC of the ribosome. Here using cryogenic electron microscopy (cryo-EM) we determined high resolution structures of the wild-type *E. coli* ribosome in complex with full-length tRNAs acylated with each of the three monomers–Apy, *o*ABZ and *m*ABZ–and accommodated within the A site of the PTC. Our structures reveal that, although the ribosome is capable of accommodating aminobenzoic acid derivatives within the A-site cleft, the size and rigid conformation of these monomers and their positioning within the PTC fails to trigger the induced-fit conformation of the ribosome and leads to loss of two of three water molecules used as proton shuttles during amide bond formation. Based on these observations, we propose a structural and mechanistic basis for the lower incorporation efficiency of aminobenzoic acid monomers relative to l-α-amino acids and each other and provide guidance on the size and geometry of non-proteinogenic monomers likely to be accepted efficiently by the wild-type PTC.

## Results

### Structures of oABZ, mABZ, and Apy within the PTC of the wild-type ribosome – map quality and monomer density

To visualize the positioning of aminobenzoic acid monomers Apy, *o*ABZ, and *m*ABZ within the ribosomal PTC, we obtained cryo-EM structures of the wild-type *E. coli* ribosome in complex with full-length acylated tRNAs in both the P and A sites. To avoid acyl-tRNA hydrolysis during sample preparation, A- and P-site monomers were appended through amide linkages using tRNA molecules carrying a 3’-amino group (3’-NH_2_-tRNA, Figure 1A, Supplementary Figures S1-S5). Purified *E. coli* ribosomes were incubated with the P-site substrate fMet-NH-tRNA^fMet^ and an A-site substrate in which 3’-NH_2_-tRNA^Phe^ was acylated with either Apy, *o*ABZ, or *m*ABZ (Figure 1A) and the structures solved using cryo-EM (Supplementary Figures S6-S8). Initial rounds of 2D and 3D classification were performed to select intact 70S ribosomal particles (Supplementary Figures S9-S11). The resulting cryo-EM maps contained complexes in the classical (non-rotated) state of the ribosome^24^ with clear density for both P- and A-site tRNAs. To improve the resolution of the Apy, *o*ABZ, and *m*ABZ monomers, we further classified the ribosomal particles based on A-site tRNA occupancy. The final maps of 70S ribosomal particles containing A-site *o*ABZ-NH-tRNA^Phe^ or Apy-NH-tRNA^Phe^ were refined to a global resolution of 1.9 Å and 2.1 Å, respectively (Supplementary Figures S6 and S7, Supplementary Tables S1 and S2). This resolution allowed for unambiguous modeling of A-site monomers *o*ABZ and Apy in a single conformation (Figure 1B, 1C). The A-site *m*ABZ-NH-tRNA^Phe^ complex with the ribosome, meanwhile, was refined to a global resolution of 2.3 Å (Supplementary Figure 8, Table S2). This resolution permitted modeling of the *m*ABZ monomer in one of two conformations related by a 180° rotation about the aryl-carbonyl bond (Figure 1B, 1C).

**Figure 1:**
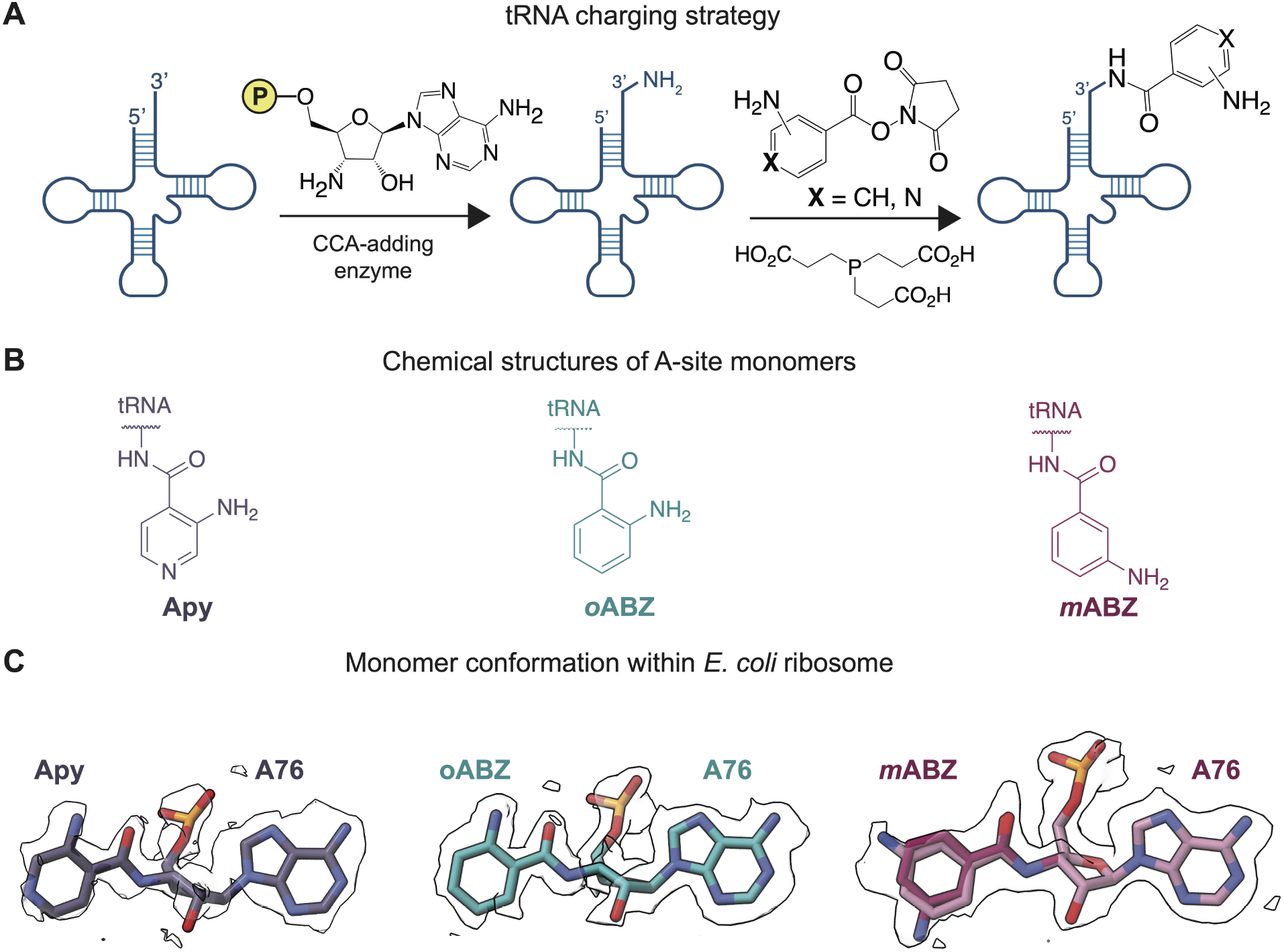
Synthesis of acylated A-site tRNAs and conformation of appended Apy, *o*ABZ, and *m*ABZ monomers within the *E. coli* PTC. (A) Strategy for preparation of 3’-NH_2_-tRNA^Phe^ acylated with aminobenzoic acid derivatives 3-aminopyridine-4-carboxylic acid (Apy), *ortho*-aminobenzoic acid (*o*ABZ), and *meta*-aminobenzoic acid (*m*ABZ). (B) Chemical structures of hydrolysis-resistant amides formed upon acylation of 3’-NH_2_-tRNA^Phe^ by Apy, *o*ABZ, and *m*ABZ. (C) Cryo-EM density showing the conformation of each aminobenzoic acid derivative within the ribosomal PTC. Maps shown were supersampled for smoothness.

### Accommodation of differentially substituted aminobenzoic acid monomers within the PTC

The A-site tRNA positions the incoming α-amino acid ester within the PTC with the aid of a base-pairing interaction between C75 of the A-site tRNA and G2553 within the A loop of the 23S rRNA.^25^ Previous structures of ribosomes containing tRNAs acylated with α-amino acids in the A and P sites reveal that when the C75-G2553 base-pairing interaction is intact, the incoming A-site α-amino acid monomer is guided into the A-site cleft, a wedge-shaped pocket created by the nucleobases of rRNA residues A2451 and C2452.^4,5,26,27^ These interactions are largely conserved when the A-site α-amino acid ester is replaced by aminobenzoic acid monomers Apy, *o*ABZ, or *m*ABZ (Figure 2). All three cryo-EM models show clear base pairing between G2553 in the A loop and C75 of the A-site tRNA and the aromatic ring of each aminobenzoic acid monomer is inserted between residues A2451 and C2452, showing near-canonical positioning compared to natural l-α-amino acids. However, because the aromatic ring is embedded within the amino acid backbone and lacks a traditional side chain, its insertion into the A2451/C2452 cleft positions the A-site tRNA ester linkage more than 1.5 Å further from the P-site monomer than the ester of a natural A-site α-amino acid (Figure 3, Supplementary Figure 12).

**Figure 2.**
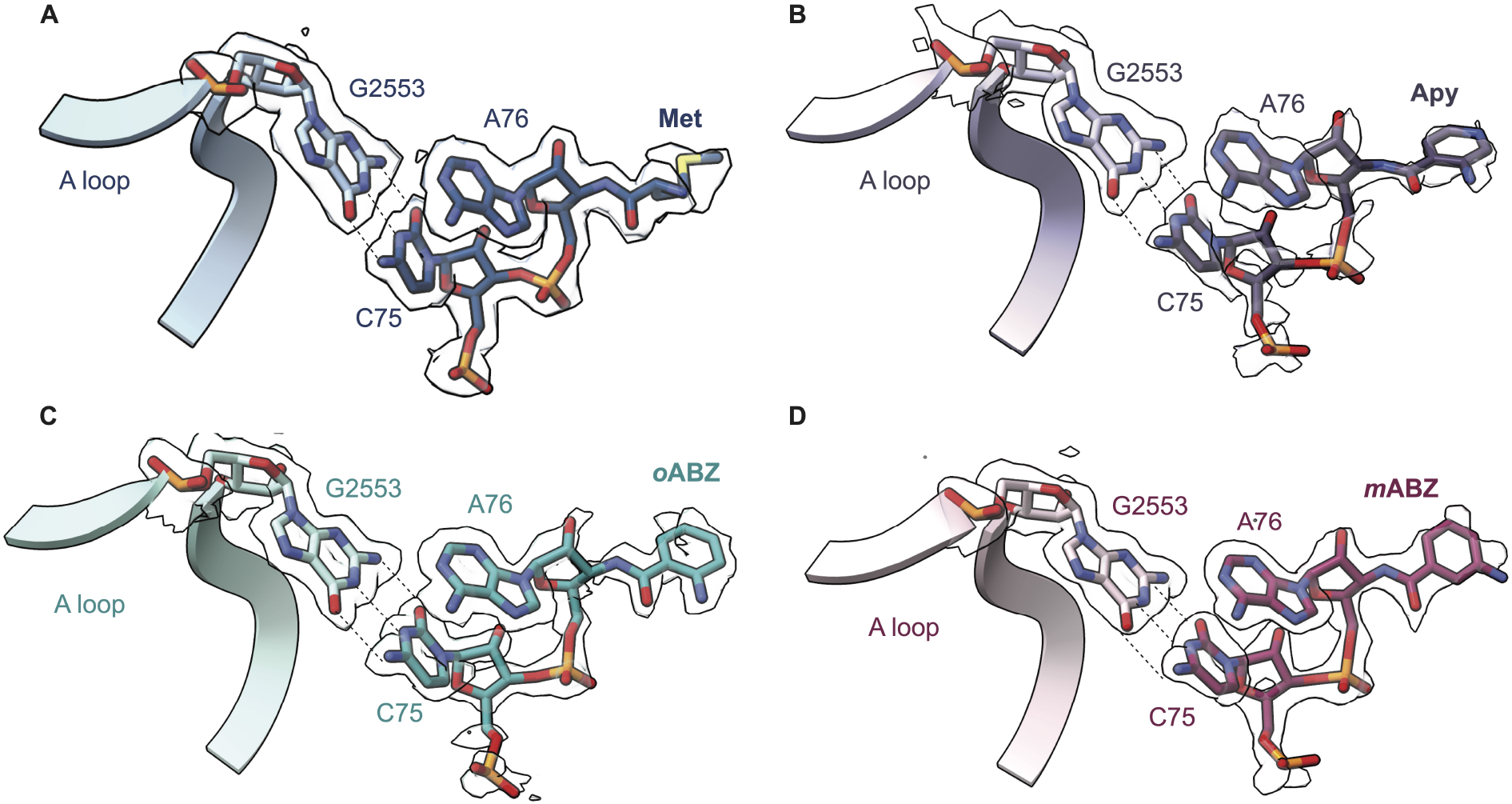
Interaction of A-site tRNAs within the PTC when acylated with (A) fMet; or aminobenzoic acid monomers (B) Apy; (C) *o*ABZ; and (D) *m*ABZ. Base pairing between C75 of the acylated A-site tRNA with G2553 within the A loop is depicted with dashed lines. Cryo-EM densities of the nucleotides and monomers are shown. Maps shown were supersampled for smoothness.

**Figure 3.**
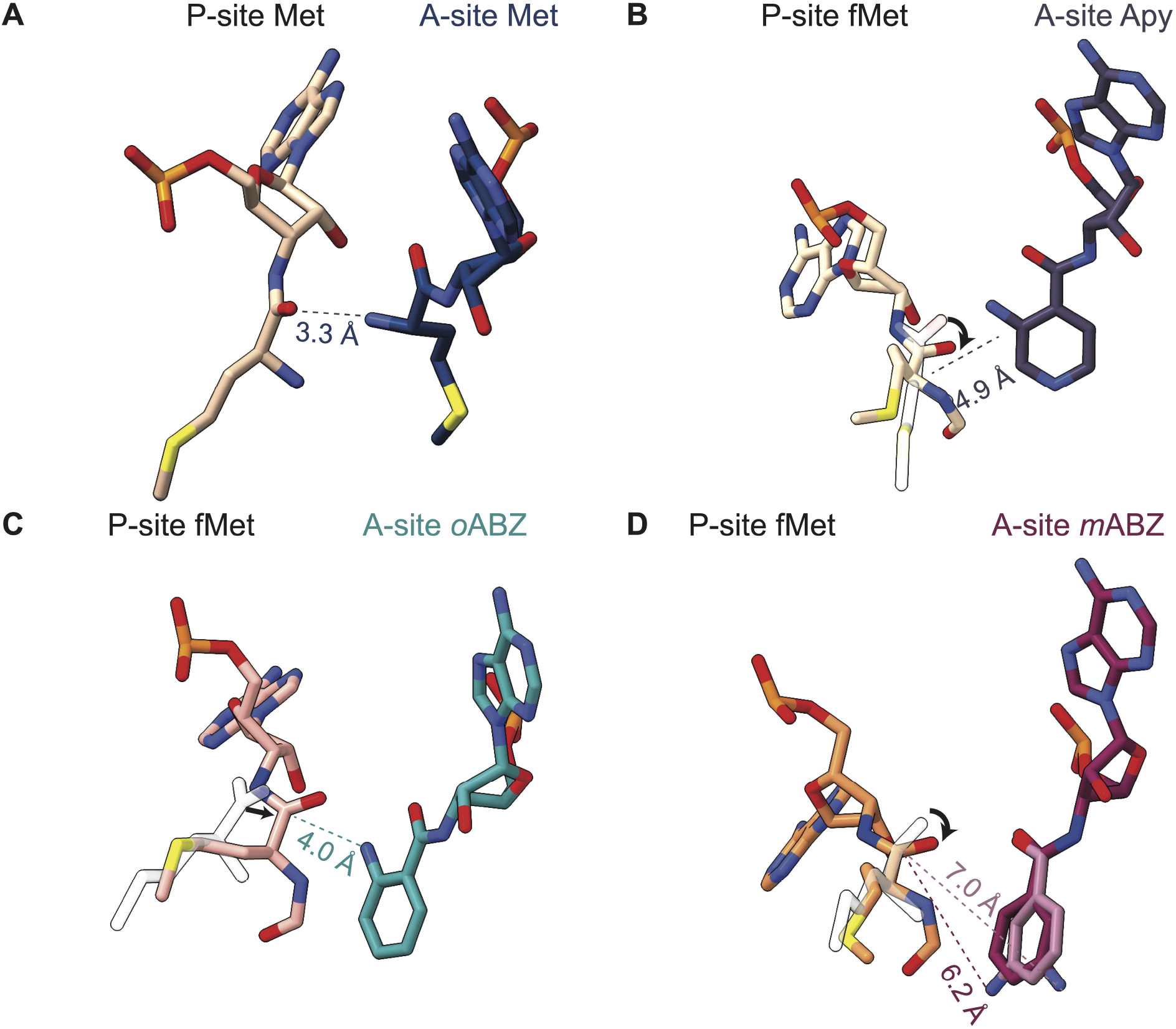
Relative positioning of the reactive groups of P- and A-site monomers. (A) P-site Met and A-site Met (B) P-site fMet and A-site Apy (C) P-site fMet and A-site *o*ABZ (D) P-site fMet and A-site *m*ABZ. The two conformations of *m*ABZ are shown in maroon and pink atomistic representation. The positioning of the carbonyl carbon and side chain of methionine from (A) are shown as transparent outlines in (B)-(D) and black arrows represent the rotation or translation of the carbonyl group.

### Positioning of the A-site amine nucleophile with respect to the P-site carbonyl electrophile

The PTC promotes peptide bond formation by positioning the nucleophilic α-amino group of the A-site aminoacyl-tRNA in proximity to the electrophilic sp^2^-hybridized carbonyl carbon of the P-site peptidyl-tRNA. Nucleophilic attack generates a tetrahedral intermediate that, after proton transfer, breaks down to form the product–a peptidyl-tRNA carrying an additional C-terminal α-amino acid. The distance between the α-amine and reactive carbonyl for a properly positioned α-amino acid is 2.9 - 3.3 Å with the expected Bürgi-Dunitz angle of 90-110° (Figure 3A).^5,16,28^ Comparison of the three cryo-EM models with aminobenzoic acid monomers in the A site revealed that, while the conserved base pair between C75 in A-site tRNA and G2553 in 23S rRNA is retained (Figure 2), the positioning of the aromatic ring in the A-site cleft formed by A2451 and C2452 shifts the position of the monomer within the PTC, especially the nucleophilic amine, relative to that of an l-α-amino acid (Supplementary Figure S12). In the case of the *m*ABZ monomer, the aromatic ring could be modeled in two distinct conformations that differ by a 180° rotation about the aryl-carbonyl bond (Figure 1C). In neither *m*ABZ conformation is the nucleophilic amine positioned to attack the P-site fMet carbonyl, as it is pointed away from the P-site fMet and separated from the reactive carbonyl by a distance of greater than 6 Å (Figure 3D). The position of the A-site *m*ABZ monomer within the PTC with respect to both distance and orientation is consistent with the low reactivity observed *in vitro*.

The high resolution of the ribosome cryo-EM maps allowed for unambiguous modeling of *o*ABZ and Apy monomers in a single conformation, with the exocyclic nucleophilic amine clearly visible (Figure 1B). In the structure containing an A-site *o*ABZ monomer, the nucleophilic amine is positioned 4.0 Å away from the P-site fMet carbonyl; in the structure containing an A-site Apy monomer, the distance is 4.9 Å (Figure 3). Notably, in all three experimental structures (Apy, *o*ABZ, and *m*ABZ), the P-site carbonyl oxygen is rotated towards the A-site monomer and lies almost in the same plane as the A-site nucleophile (Figure 3).^5,16,28^. Although these distances and orientations differ from those seen for properly positioned α-amino acids,^5,16,23^ the observation of *in vitro* reactivity suggests that, at least for Apy and *o*ABZ, the exocyclic amino group is positioned just close enough to allow amide bond formation under physiological conditions. Taken together, the orientation and elongated distance between attacking amine and carbonyl carbon seen when the A-site contains *m*ABZ versus Apy, *o*ABZ or a natural α-amino acid is consistent with the diminished reactivity of *m*ABZ in *in vitro* translation reactions. However, the structures alone do not explain the higher reactivity of Apy relative to *o*ABZ. The only structural difference we detect in the models is a slightly more diffuse density surrounding the P-site fMet and A76 ribose when the ribosome A-site is occupied by Apy, suggesting increased flexibility of the P-site monomer (Supplementary Figure S12). It is possible that this increased flexibility is one contribution to the increased reactivity of Apy compared to *o*ABZ (*vide infra*).

### Alternative positioning of inner shell nucleotides in the PTC

Nucleotide U2506 in the PTC has been implicated in the induced fit mechanism necessary for efficient amide bond formation by the ribosome. In the uninduced state, when the A site is not occupied by a natural aminoacyl-tRNA, U2506 forms a wobble base pair with G2583.^4,29^ Once an aminoacyl-tRNA is accommodated in the A site, the ribosome transitions to an induced state in which U2506 is re-positioned to confine the side chain of the A-site α-amino acid within the A-site cleft and away from the reaction center (Figure 4A). U2506 repositioning also helps position the nucleophilic α-amine within ~3 Å of the P-site carbonyl carbon.^4,29^ This repositioning is not observed in structures containing aminobenzoic acid monomers. In these cases, docking of the conformationally restricted aromatic ring in the A-site cleft between A2451 and C2452 causes the ring to abut the backbone of U2506. This steric restriction forces U2506 to rotate by ~70° away from the position it occupies in the induced state seen in ribosomes with natural α-amino acid monomers in the A site (Figures 4B-D). The steric restriction also moves the exocyclic O2 of the U nucleobase about 6.8 Å away from the Cα of the A-site monomer, compared a distance of 4 Å in case of an A-site methionine (Supplementary Figure S13).^16^ In addition, U2585, which normally moves in tandem with U2506 in the induced fit mechanism to expose the P-site carbonyl for attack, remains in the uninduced position within van der Waals distance of the P-site fMet (Figure 5A, 5B). This close positioning of U2585 to the acyl bond rotates the carbonyl out of the optimal position for nucleophilic attack, consistent with the conformation seen in the uninduced state (Figure 5A, 5D-5F).^4,29^ Moreover, A2602, which coordinates one of the three waters that comprise the “proton wire” mechanism for amide bond formation^5,30–32^, is disordered in the three ribosome structures and could not be modeled with high confidence (Supplementary Figure S14). Taken together, the structures of ribosomes containing aminobenzoic acid monomers in the A site all adopt conformations closer to the uninduced state of the PTC, rather than closing around the substrates and helping position water molecules for catalysis, as observed with l-α-amino acids.^5,30–32^

**Figure 4:**
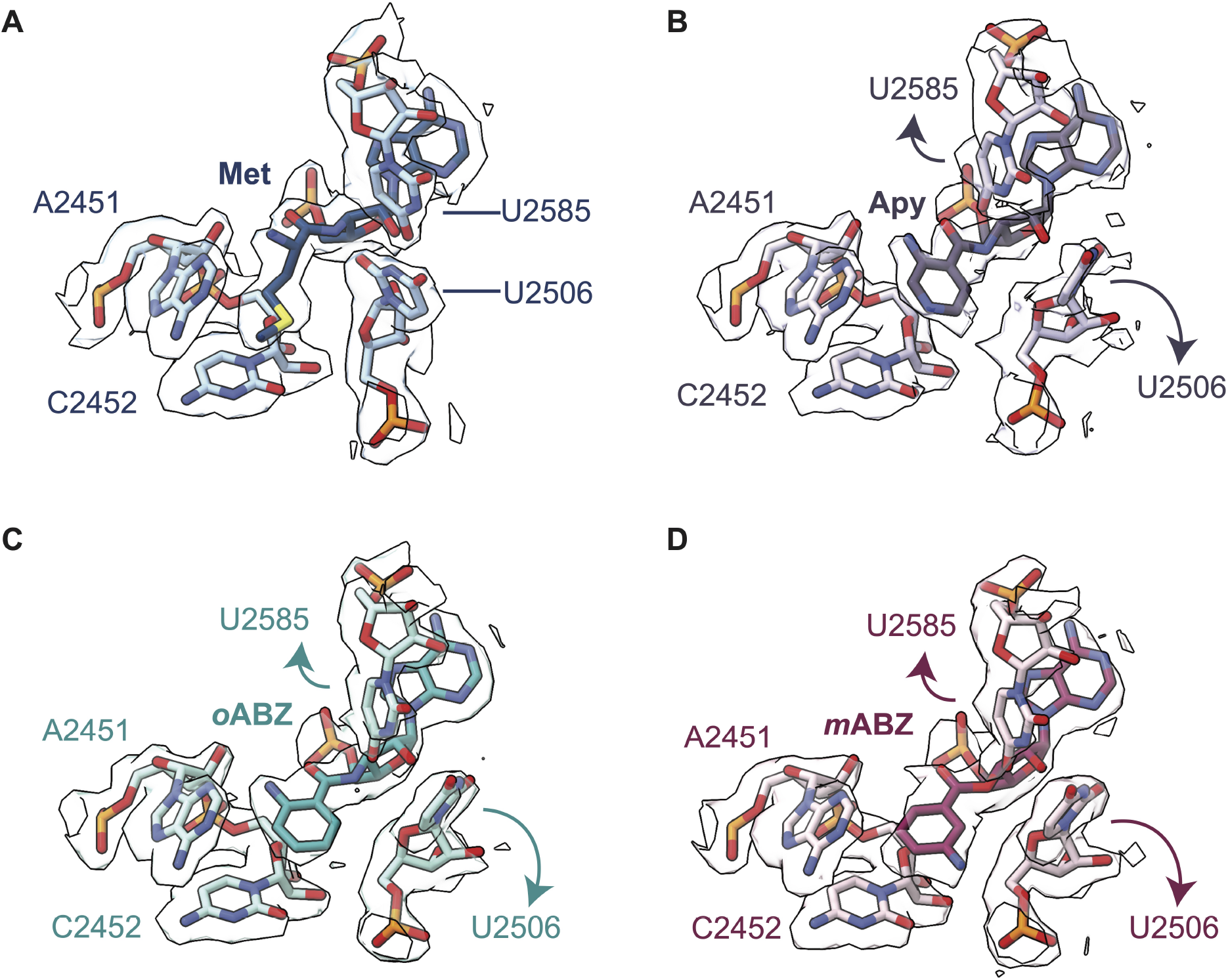
Aminobenzoic acid monomers hinder A-site tRNA-promoted induced fit within the ribosomal PTC. (A) Close-up of the PTC illustrating how U2506 encloses the side chain of the natural l-α-amino acid methionine within the A-site cleft formed by nucleotides A2451 and C2452. (B-D) This induced conformational change is not observed when the A-site cleft is occupied by aminobenzoic acid monomers Apy (B); *o*ABZ (C); or *m*ABZ (D). In these cases, the aminobenzoic acid monomer sterically impedes the movement of U2506 and prevents it from adopting the position seen when the A-site is occupied by a canonical α-amino acid. As a result, U2506 along with U2585 cannot adopt the induced conformation required for rapid bond formation. Cryo-EM density for nucleotides U2506 and U2585 are shown as outlines. Maps were supersampled for smoothness.

**Figure 5:**
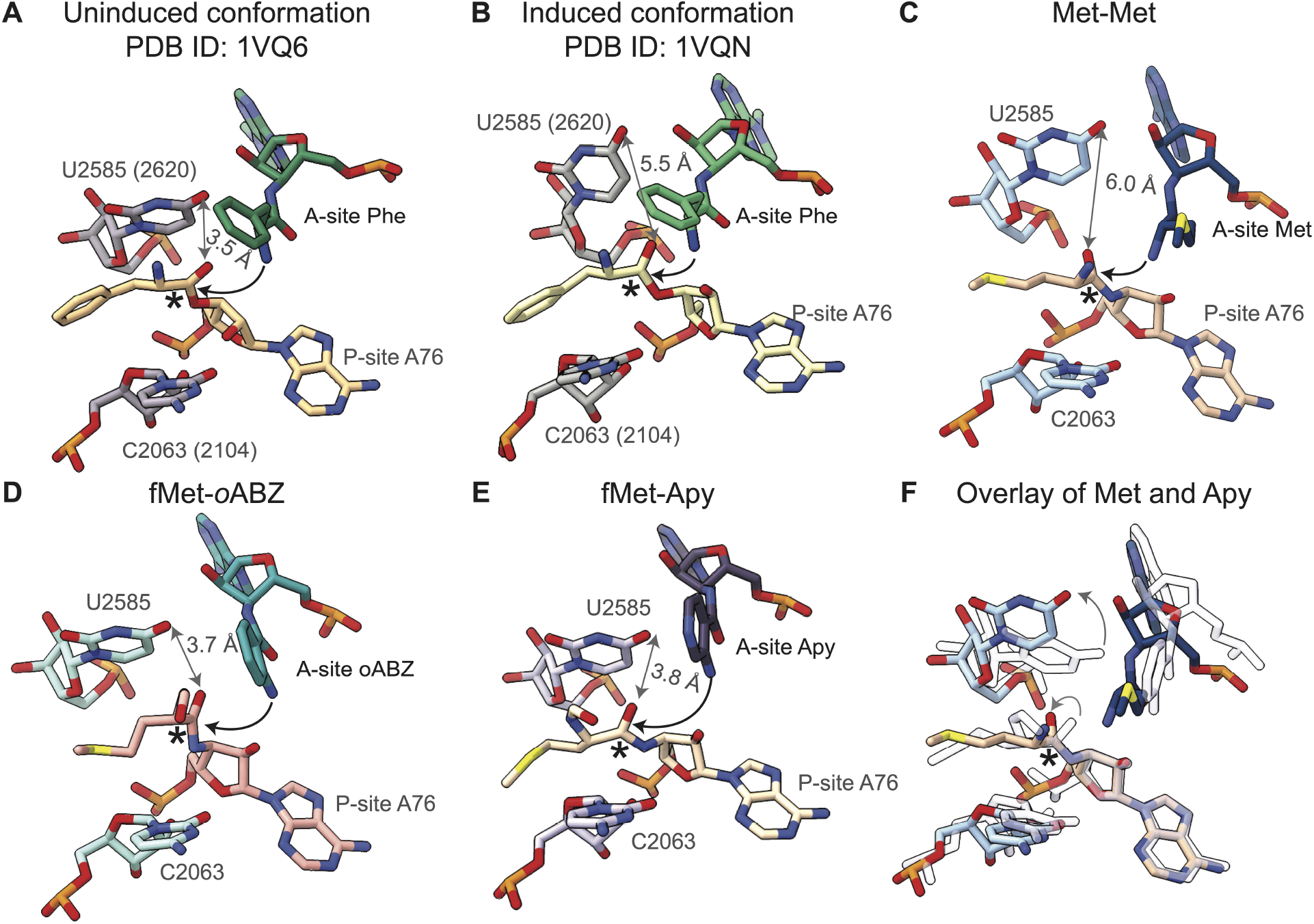
Rigid conformation of aminobenzoic acid monomers inhibits conformational changes associated with induced fit of A- and P-site tRNAs within the PTC. (A) The uninduced conformation of the PTC (PDB ID 1VQ6), in which nucleotides U2585 and C2063 protect the P-site carbonyl (denoted by an asterisk) from nucleophilic attack. 23S numbering of *Haloarcula marismortui,* as used in these crystal structures, are indicated in parentheses and the distance between O6 on U2585 and P-site carbonyl oxygen indicated in grey. Black arrows represent the attack of A-site nucleophile on P-site electrophile. (B) The induced conformation observed when a natural acyl-tRNA occupies the A-site (PDB ID 1VQN). When this substrate is bound properly, U2585 moves to expose the P-site carbonyl group to the A-site nucleophile. (C) The structure of the PTC with the natural α-amino acid, Met, in the P and A sites resembles the induced conformation seen in PDB ID 1VQN. (D, E) The structure of the PTC with (D) *o*ABZ and (E) Apy aminobenzoic acid monomers in the A-site show U2585 in a position that resembles the un-induced conformation. In these two structures, U2585 is within van der Waals contact of the P-site fMet, preventing its carbonyl group from adopting a reactive conformation. (F) Overlay showing differences in the position of U2585 in structures containing Met and Apy monomers (as a representative example) in the A site. The structure with the natural α-amino acid, Met, is shown in shades of blue, whereas the structure with Apy is shown with outlines.

## Discussion

### Cryo-EM structures reveal positioning of aminobenzoic acid derivative monomers in the PTC

The cryo-EM structures reported here represent the first high-resolution maps of ribosomes whose active sites contain monomers that are not α-amino acids. Advances in cryo-EM, along with the use of hydrolytically stable acyl-tRNAs combine to produce structural models of ribosome complexes characterized by unprecedented levels of detail. In particular, as all three aminobenzoic acid monomers contain aromatic rings that constrain their conformational freedom, their positions within the PTC are extremely well-ordered. In the case of *o*ABZ, where the final cryo-EM map reached a global resolution of 1.9 Å, a slight torsion (~12°) of the aromatic ring relative to the carbonyl group was evident. The global resolution for the complexes containing Apy and *m*ABZ were slightly lower (2.0 Å and 2.3 Å respectively), yet in both cases the aromatic ring and exocyclic amino substituent are clearly visible. The high resolution also revealed *m*ABZ to be bound in two different conformations rotated by 180°, in both cases with the nucleophilic amine pointed away from the P-site monomer (Figure 3D).

Structures of the ribosome with natural l-α-amino acids show that the A-site amine nucleophile is typically located at a distance of about 3 Å away from the carbonyl carbon of the P-site amino acid and oriented for nucleophilic attack.^5,16^ Recently, metadynamics simulations suggest that monomers with nucleophile-carbonyl distances of up to 4 Å in the ground state are likely to be reactive within the PTC.^16^ In the present structures, we observe that the nucleophilic amine of *m*ABZ is located greater than 6 Å away from the carbonyl carbon of the P-site amino acid, consistent with the extremely low reactivity of this monomer *in vitro*. In the cryo-EM structure with *o*ABZ, the amine to carbonyl distance is 4 Å, a distance right at the cusp of the metadynamics prediction for reactive monomers. Thus, based on the distances alone, the intermediate incorporation efficiency of *o*ABZ relative to *m*ABZ and a natural α--amino acid can be rationalized, and is consistent with the metadynamics simulations. However, Apy poses a conundrum. It is incorporated into peptides more efficiently than *o*ABZ, although the cryo-EM structure shows that the spacing between the reactive groups is greater, at about 5 Å. The additional observation that the P-site fMet and the A76 ribose appears to be more diffuse in the Apy structure compared to the *o*ABZ structure may reflect increased dynamics in the PTC that leads to increased Apy reactivity.

### Rigid conformation of aminobenzoic acid derivative monomers inhibits conformational changes associated with the induced fit mechanism

The high-resolution structures reported here also identify a second explanation for the diminished reactivity of aminobenzoic acid monomers in *in vitro* translation reactions. In each case, the expanded and conformationally rigid aromatic backbone sterically blocks the induced fit of the A- and P-site substrates that occurs when natural acyl-tRNAs engage within the A site. Upon binding of a natural acyl-tRNA, nucleotide U2506 rotates by 90° and confines the α-amino acid side chain within the A-site cleft and away from the reaction center, while also positioning the carbonyl carbon of the A-site α-amino acid close to the 2’ OH of A76 of the P-site tRNA (Figure 5A, 5B). The end result is a conformation in which the α-amine of the A-site acyl-tRNA is positioned optimally for amide bond formation.^4,5,16,28,29^ As observed in all three structures presented here, the aromatic rings of the monomers pack tightly against the backbone of nucleotide U2506 in 23S rRNA, part of the inner ring of nucleotides in the PTC (Figure 4, Supplementary Figure S13). This packing prevents U2506 from adopting the position seen with natural acyl-tRNA substrates and thus fails to position the attacking nucleophile with respect to the carbonyl carbon of the P-site monomer.

The inability of U2506 to rotate into the induced conformation likely also prevents movement of U2585 away from the carbonyl carbon of the P-site ester linkage, which in turn prevents its rotation into a position suitable for efficient amide bond formation (Figure 4, Figure 5). In the uninduced conformation, U2585 is within van der Waals contact distance of the P-site ester, which causes the carbonyl group to point towards the A site in an orientation unsuitable for nucleophilic attack (Figure 5A). In this conformation U2585 protects the P-site ester carbonyl from premature hydrolysis in the absence of an A-site monomer. In the induced fit mechanism, correct binding of the A-site monomer causes U2506 to rotate 90°, as described above, which in turn shifts U2585 away from the P-site ester, allowing it to rotate and expose itself to nucleophilic attack by the A-site monomer (Figure 5B). In the case of the aminobenzoic acid derivatives, the inability of the monomers to induce the conformational change in U2506 results in U2585 remaining in a position to block the P-site carbonyl carbon from rotating into an orientation suitable for nucleophilic attack (Figure 5D-F). The obstruction of the U2506 and U2585 rearrangements necessary for the induced fit mechanism, together with the increased nucleophile-carbonyl distances in the case of the aminobenzoic acid derivatives, explains the overall lower reactivity of these monomers compared to natural l-α-amino acids. The absence of the conformational changes associated with the induced fit mechanism provide a second explanation for the diminished reactivity of aminobenzoic acid monomers in *in vitro* translation assays.

The lack of conformational changes associated with the induced fit mechanism are also seen in structures of the ribosome with stalling peptides. In a cryo-EM structure of SecM in the WT *E. coli* ribosome, insertion of residue R163 in the stalling sequence into the A-site cleft PTC causes U2506 and U2585 to adopt conformations consistent with the uninduced state, resulting in translational stalling.^33^ In the case of *m*ABZ, these factors, combined with the fact that the nucleophilic amine is pointed away from the P-site render this monomer virtually unreactive. The directionality of the nucleophilic amine is reminiscent of the case of D-α-phenylalanine, for which a recent crystal structure revealed that the A-site amino group is also pointed away from the P-site carbonyl carbon resulting in the lower reactivity of this enantiomer despite the ribosome PTC adopting the induced state.^26^ On the other hand, although the inability of the ribosome to adopt the induced conformation in the presence of both Apy and *o*ABZ helps to explain their lower reactivity compared to l-α-amino acids, the reason for the reactivity difference between Apy and *o*ABZ remains more subtle. In the cryo-EM maps, the P-site fMet in the Apy complex appears to be more dynamic than in the *o*ABZ complex. With the increased dynamics suggested by the Apy ribosome complex, the P-site monomer may possess enough conformational flexibility to adopt a position compatible with nucleophilic attack, as opposed to the P-site fMet in the case of *o*ABZ, which appears to be well ordered and trapped in the uninduced conformation.

### Relative reactivity of aminobenzoic acid derivatives

Amide bond formation within the PTC requires more than the attack of a nucleophilic amine on a proximal ester carbonyl. A key consideration is the shuttling of protons between the attacking and leaving groups involved in the reaction. In the case of natural α-amino acid esters, the nucleophilic amine in the A-site exists in the ammonium form at physiological pH and must be deprotonated prior to nucleophilic attack on the P-site carbonyl carbon to form a tetrahedral intermediate. Following this step, this amine must be deprotonated once again, and a proton must be transferred to the 3’-oxygen leaving group of the P-site A76 to facilitate breakdown of the tetrahedral intermediate. There is significant evidence that these proton transfers are mediated by a trio of water molecules, referred to as W1, W2, and W3, that form a proton wire (Supplementary Figure S14).^5^ Proton shuttle mechanisms characterized by distinct 8- and 6-membered rings have also been proposed.^5,30–32^

The inability of the PTC to adopt an induced-fit conformation with aminobenzoic acid monomers bound in the A site disrupts the water positions associated with the proton wire model. For instance, nucleotide A2602, which coordinates both W1 and W2 of the proton wire, is disordered and could not be modeled with high confidence. Also, no density for the N-terminus of uL27 is seen in these structures. As a result, W1, which is proposed to act twice as a general base–first to indirectly deprotonate the attacking amine and again to deprotonate the nitrogen of the tetrahedral intermediate–is not observed. Density corresponding to W2, the proposed general acid that stabilizes the oxyanion of the tetrahedral intermediate, is also absent in these structures. The only water observed with confidence is W3, which facilitates breakdown of the tetrahedral intermediate (Supplementary Figure S14).

Why then do Apy and *o*ABZ participate in amide bond formation within the ribosome PTC? We propose that these monomers may react through a mechanism that does not require W1 or W2. The *pK*_a_ values of the acidic (ammonium) forms of the *o*ABZ and Apy methyl esters were determined to be 2.8 and −3.0, respectively, using computation (Jaguar DFT, Schrödinger Maestro Suite, release 2022-4).The *pK*_a_ value of the acidic (ammonium) form of *m*ABZ was found to be 3.5. These low *pK*_a_ values imply that none of the aminobenzoic acid monomers require the assistance of W1 as a general base prior to or subsequent to nucleophilic attack. They are expected to exist predominantly in the neutral form at neutral pH, even after the tetrahedral intermediate forms. Thus, the easily-lost proton on the amine after nucleophilic attack could compensate for the absence of W2. Given this loss would be easier from the Apy intermediate due to its lower *pK*_a_ value (−3.0 *versus* 2.8), this provides a potential explanation for the increased reactivity of Apy compared to *o*ABZ.

### Implications for the design of unnatural monomers

The incorporation of even a single unnatural monomer into a peptide chain necessitates the successful execution of several steps, beginning with charging of tRNAs with the monomers and their incorporation into elongated polymer chains. While the structures presented here reveal the details of how the aminobenzoic acid derivatives are positioned within the PTC and their inability to trigger the induced state of the ribosome necessary for efficient amide bond formation, they only provide insight into the possible defects in that single step. Other factors, such as translocation of the growing polymer chain from the A site to the P site, positioning of the monomer incorporated in the growing polymer within the P site, and interactions within the exit tunnel remain unknown. Furthermore, factors affecting other aspects of translation, such as efficiency of charging monomers onto tRNA, editing activity of mischarged tRNAs by aminoacyl-tRNA synthetases,^34,35^ and efficiency of delivery of charged tRNAs to the ribosome by EF-Tu,^36^ would also influence overall yields of the final polymer. Nevertheless, the structures of the aramid monomers within the A site of the ribosome presented here provide valuable insights into the requirements for efficient monomer incorporation by the ribosome. For example, monomers related to *o*ABZ and Apy but possessing an expanded or alternative aromatic core may allow the monomer to enter the A-site cleft without steric constraints with U2506. Monomers could be chosen to allow U2506 to rotate into its correct position in the induced fit mechanism, thereby triggering movement of U2585 and the other conformational rearrangements necessary for amide bond formation to take place efficiently. Alternatively, the conformational rearrangements associated with induced fit could be promoted using ribosome evolution strategies that exploit recently mapped intra-ribosome allosteric relationships to avoid universally conserved nucleotides.^37^ Regardless, the three ribosome models reported here provide an unprecedented structural and mechanistic rationale for differences in reactivity among non-l-α-amino acid monomers and identify clear stereochemical constraints on the size and geometry of non-proteinogenic monomers that are likely to be processed efficiently by wild-type ribosomes.

## Supporting information

Maps and Models in PTC region

## Acknowledgments

This work was supported by the NSF Center for Genetically Encoded Materials (C-GEM), CHE 2002182. We would also like to thank Dr. Dan Toso, Dr. Ravindra Thakkar, and Paul Tobias for help with cryo-EM data collection and management, and Amos Nissley for helpful comments on the manuscript. We would like to thank Dr. Fred Ward for help with protein purification. A.S. is a Chan Zuckerberg Investigator.

## Data Availability

Atomic coordinates have been deposited with the Protein Data Bank under accession codes ___, ___, and ___ (for *m*ABZ, *o*ABZ, and Apy, respectively). Cryo-EM maps have been deposited with the Electron Microscopy Data Bank under the accession codes EMDB-___ (70S global map for *m*ABZ) and EMDB- ___(50S focused refinement map *m*ABZ); EMDB- ___(70S global map for Apy) and EMDB- ___(50S focused refinement map *m*ABZ); EMDB-___(70S global map for *o*ABZ), EMDB- ___(50S focused refinement map *o*ABZ), and EMDB- ___(30S focused refinement map *o*ABZ).

## Supporting Information

The supporting information is available free of charge at ____.

- Materials and methods, cryo-EM data collection and model refinement statistics, NMR characterization of aminobenzoic acid derivatives, supplementary figures.

