## Supplementary Data for "Aminobenzoic acid derivatives obstruct induced fit in the catalytic center of the ribosome"

**Table of Contents**


1. Materials and Methods
    a. Purification of E. coli CCA adding enzyme
    b. Purification of E. coli methionyl-tRNA-formyltransferase (MTF)
    c. Transcription of $tRNA^{fMet}$(-A) and $tRNA^{Phe}$ (-A)
    d. Preparation of tRNA molecules carrying a 3'-amino group, 3'-$NH_2$-$tRNA^{fMet}$ and 3'-$NH_2$-$tRNA^{Phe}$
    e. Enzymatic acylation of 3'-$NH_2$-$tRNA^{fMet}$ with fMet
    f. Synthesis of 3-azidopyridine-4-carboxylic acid-N-hydroxysuccinimide (NHS) ester
    g. Synthesis of 2-azidobenzoic acid
    h. Synthesis of 2-azidobenzoic acid-N-hydroxysuccinimide (NHS) ester
    i. Synthesis of 3-azidobenzoic acid
    j. Synthesis of 3-azidobenzoate-N-hydroxysuccinimide (NHS) ester
    k. Acylation of 3'-$NH_2$-$tRNA^{Phe}$
    l. Purification of Acylated 3'-$NH_2$-$tRNA^{Phe}$
    m. Cryo-EM complex preparation
    n. Cryo-EM sample preparation
    o. Cryo-EM data collection
    p. Image Processing
    q. Pixel Size Calibration
    r. Modeling
    s. pKa calculations
2. Supplementary tables
    a. Table S1: Data collection and processing for WT *E. coli* ribosome complexes with aminobenzoic acid monomers
    b. Table S2: Model Refinement Statistics for WT *E. coli* ribosome complexes with aminobenzoic acid monomers.
    c. Table S3: Sequences of RNA oligomers used in this study
    d. Table S4: Sequences of DNA primers used in this study. mG represents 2'-O-methylguanosine modification

3. Supplementary figures
    a. Figures S1-S5: Synthesis, tRNA acylation and characterization of aminobenzoic acid derivatives
    b. Figures S6-S8: Resolution estimates of cryo-EM maps
    c. Figures S9-S11: Cryo-EM data processing workflow
    d. Figure S12: Relative positioning of the reactive groups of P- and A-site monomers showing cryo-EM density of the P- and A-site monomers.
    e. Figure S13: Repositioning of nucleotide U2506 in the complexes containing aminobenzoic acid derivatives.
    f. Figure S14: Binding of aminobenzoic acid monomers in the A-site disrupts the proton wire proposed to support proton transfers during amide bond formation
    g. Figures S15-S26: $^{1}H$ and $^{13}C$ NMR characterization of of compounds 1-6
4. References

---

**Materials and Methods**

*Purification of E. coli CCA adding enzyme*

Bacterial glycerol stocks transformed with an expression plasmid encoding the *E. coli* CCA adding enzyme (CTP(ATP): tRNA nucleotidyl transferase) was a gift from Prof. Yuri Polikanov (University of Illinois Chicago, Chicago USA). Recombinant His-tagged *E. coli* CCA adding enzyme was purified by adapting protocols used previously.[1,2] Briefly, two 1 L cultures of Luria broth (LB) supplemented with 34 µg/mL chloramphenicol were inoculated with a 1:20 dilution of an overnight culture (grown at 37 ºC) and incubated at 37 ºC, 180 rpm until they reached an $OD_{600}$=1.0. Protein overexpression was induced with 250 µM Isopropyl-β -D-1-thiogalactopyranoside (IPTG), and the cultures were grown for an additional 3 hours at 37 ºC. The cells were pelleted for 30 minutes at 4650 ×g (F9-6X1000LEX rotor, Thermo Scientific), resuspended in 30 mL of Buffer A (50 mM Tris-HCl pH 8.0, 300 mM NaCl, 5%

glycerol (%v/v), 2 mM imidazole), and stored at -80 ºC. The pellets were subsequently thawed in an ice-water slurry for 30 minutes, lysed by sonication, and the lysate clarified by centrifugation at 9280 ×g for 30 minutes (F14-14X50CY rotor, Thermo Scientific). The supernatant was loaded onto a 5 mL HisTrap column (Cytiva) pre-equilibrated with Buffer A and the flowthrough was recirculated over the column for 1 hour to ensure maximum binding of the enzyme. The bound protein was washed with 15 column volumes (CV) of Buffer A, and then eluted using a linear gradient of 0-100% Buffer B (50 mM Tris-HCl pH 8.0, 300 mM NaCl, 5% glycerol (%v/v), 300 mM imidazole). Fractions containing CCA-adding enzyme (Molecular weight (MW) 48 kDa, analyzed on a 12% SDS-PAGE gel) were pooled and concentrated to 2 mL and then loaded onto a Superdex pg 75 16/600 column (GE Healthcare) pre-equilibrated with GF buffer (40 mM Tris-HCl pH 8.0, 0.5 mM EDTA, 5% glycerol (%v/v), 7 mM β-mercaptoethanol), and eluted over 1.5 CV. Fractions were analyzed using a 12% SDS-PAGE gel and pure fractions were pooled, concentrated, flash frozen in liquid nitrogen, and stored as single use aliquots. The final concentration was estimated to be 91 µM using the $A_{280}$ absorbance ($\varepsilon_{280}$ = 55,920 $M^{-1}cm^{-1}$)[3].

*Purification of E. coli methionyl-tRNA-formyltransferase (MTF)*

The expression plasmid for *E. coli* MTF was a gift from Patrick Ginther, Schepartz lab, and was transformed into BL21(DE3) cells for overexpression. Two cultures of 1 L LB supplemented with 100 µg/mL were inoculated with a 1:20 dilution of overnight cultures (grown at 37 ºC) and incubated at 37 ºC, 150 rpm till they reached an $OD_{600}$ = 0.4. Protein overexpression was induced with 500 µM IPTG and incubation was continued overnight at 16 ºC. Cells were pelleted 30 minutes at 4650 ×g (F9-6X1000LEX rotor, Thermo Scientific), and resuspended in 20 mL Wash Buffer (20 mM HEPES-KOH pH 7.5, 150 mM NaCl, 10 mM $MgCl_2$, 1 mM DTT, 20 mM imidazole). The cells were then lysed by sonication and the lysate clarified by centrifugation at 25,000 ×g (JA-20 rotor, Beckman Coulter). The supernatant was loaded onto a 1 mL HisTrap column (GE Healthcare) pre-equilibrated with Wash Buffer, and the flowthrough was recirculated over the column for 1 hour to ensure maximum binding of the enzyme. The bound protein was washed with 5 CV of Wash Buffer, and then eluted using a linear gradient of 0-100% Elution Buffer (20 mM HEPES-KOH pH 7.5, 150 mM NaCl, 10 mM $MgCl_2$, 1 mM DTT, 500 mM imidazole). Fractions containing MTF (MW 30 kDa, analyzed on a 12% SDS-PAGE gel) were pooled and dialyzed overnight at 4 ºC against Storage Buffer (20 mM HEPES pH 7.5, 150 mM NaCl, 10 mM $MgCl_2$, 1 mM DTT, 20% (v/v) glycerol) with a 7K molecular weight cut-off (MWCO) cassette. The protein was concentrated to 1.5 mL and final concentration was estimated to be 263 µM using the $A_{280}$ absorbance ($\varepsilon_{280}$ = 44,460 $M^{-1}cm^{-1}$).[3]

*Transcription of tRNA$^{fMet}$(-A) and tRNA$^{Phe}$ (-A)*

Using previously described protocols, DNA templates for *E. coli* tRNA$^{fMet}$(-A) with a C1G mutation (to improve transcription yield) and tRNA$^{Phe}$(-A) (Table S3) were prepared using a double PCR amplification method described previously.[4–6] Briefly, long overlapping primers, fMet_C1G_temp_Fw and fMet_C1G_temp_Rv(-A) for tRNA$^{fMet}$(-A), or Phe_temp_Fw and Phe_temp_Rv for tRNA$^{Phe}$(-A) (Table S4) were PCR amplified using Q5 polymerase (NEB). The PCR products were agarose gel purified and amplified using short primers fMet_C1G_amp_Fw and fMet_C1G_amp_Rv(-A) for tRNA$^{fMet}$ and Phe_amp_Fw and Phe_amp_Rv(-A) for tRNA$^{Phe}$ using Q5 DNA polymerase (NEB). Amplification reactions were extracted using phenol-chloroform, and precipitated in EtOH. The amplified PCR products were quantified using an agarose gel against a known standard (NEB Quick-Load 1 kb Plus DNA ladder). tRNA$^{fMet}$(-A) and tRNA$^{Phe}$(-A) were *in vitro* transcribed in transcription buffer (50 mM Tris-HCl pH 7.5, 15 mM $MgCl_2$, 5 mM DTT, 2 mM spermidine), 2.5 mM NTPs (Promega), 1 U/µL murine RNase inhibitor (NEB), 0.5 U/µL T7 RNA polymerase (NEB), 0.5 U/µL YIPP (NEB), and 50 ng/µL DNA template. Reactions were incubated at 37 ºC for 16 hours following which DNA templates were digested with 45 U/mL RQ1 DNase (Promega) at 37 ºC for 30 minutes. The tRNAs were then precipitated using EtOH and purified using a 12% polyacrylamide, 7 M urea, 1X TBE (100 mM Tris, 100 mM boric acid, 2 mM EDTA disodium salt) gel. Gel slices were excised and tRNAs extracted by crushing and soaking the gel in 300 mM NaOAc pH 5.2 overnight. The tRNAs were then precipitated in EtOH, the pellets dried, and then resuspended in nuclease-free water prior to storage at -80 ºC.[4–6]

*Preparation of tRNA molecules carrying a 3'-amino group, 3'-$NH_2$-tRNA$^{fMet}$ and 3'-$NH_2$-tRNA$^{Phe}$*

tRNA$^{fMet}$(-A) and tRNA$^{Phe}$(-A) were converted into 3'-$NH_2$-tRNA$^{fMet}$ and 3'-$NH_2$-tRNA$^{Phe}$ using 2'-amino-2'-deoxyadenosine-5'-triphosphate (Axxora) and either *A. fulgidus* CCA adding enzyme (gift from Fred Ward, Cate lab) or *E. coli* CCA adding enzyme using previously described protocols.[4–6] Successful elongation to generate 3'-$NH_2$-tRNA$^{fMet}$ and 3'-$NH_2$-tRNA$^{Phe}$ was confirmed by observing a gel shift on a 12% polyacrylamide/7M urea/1X TBE gel.

*Enzymatic acylation of 3'-$NH_2$-tRNA$^{fMet}$ with fMet*

3'-$NH_2$-tRNA$^{fMet}$ was enzymatically aminoacylated using methionyl-tRNA synthetase (MetRS)[5,6] and methionyl-tRNA formyltransferase (MTF) (gifts from Fred Ward, Cate lab). 10 µM 3'-$NH_2$-tRNA$^{fMet}$ was incubated with 1 µM MetRS and 1 µM MTF in aminoacylation buffer (50 mM HEPES pH 7.5, 20 mM $MgCl_2$, 10 mM KCl, 2 mM dithiothreitol (DTT), 10 mM ATP) along with 1:40 volume murine RNase inhibitor (NEB), 10 mM methionine, and 300 µM 10-formyltetrahydrofolate.[7] The reactions were incubated at 37 ºC for 30 minutes, extracted with phenol chloroform, and precipitated with ethanol. Charging was confirmed by observing a gel shift on a 12% polyacrylamide/7M urea/1X TBE gel. The charged fMet-NH-tRNA$^{fMet}$ was stored in nuclease free water at -80 ºC prior to use.

*Synthesis of 3-azidopyridine-4-carboxylic acid (1):* The synthesis of **(1)** was adapted from previous reports (Supplementary Figures S1, S15-16).[8] 2-aminopyridine-4-carboxylic acid (Apy, 1 equivalency, 504 mg, 3.65 mmols) was dissolved in 10 mL of 10% HCl and stirred on ice for 10 minutes to form a partially insoluble suspension. Sodium nitrite (1.2 equivalencies, 302 mg, 4.38 mmols) was dissolved in 1.5 mL of water and slowly added to the suspension of 2-aminopyridine-4-carboxylic acid. The resulting solution was allowed to stir at room temperature for 15 minutes. Then, sodium azide (1.2 equivalencies, 285 mg, 4.38 mmols) was dissolved in 1 mL of water and carefully added to the solution. The reaction was stirred at room temperature for 1 hour. The reaction mixture was transferred to a separatory funnel and the aqueous phase was extracted with 20 mL of dichloromethane (3x) and 20 mL of ethyl acetate (3x). The combined organic extracts were dried over sodium sulfate and dried *in vacuo* to yield **(1)** as a semi-beige solid in 30% yield (178 mg, 1.1 mmols). $^1$H NMR (500 MHz, DMSO-$d_6$): δ 13.83 (s, 1H), 8.67 (s, 1H), 8.48 (d, 1H, J = 5.0 Hz), 7.66 (d, 1H, J = 5.0 Hz). $^{13}$C NMR (500 MHz, DMSO-$d_6$): δ 165.37, 146.00, 143.63, 134.26, 130.56, 123.37.

*Synthesis of 3-azidopyridine-4-carboxylic acid-N-hydroxysuccinimide (NHS) ester (2):* The synthesis of (**2**) was adapted from previous reports (Supplementary Figures S1, S17-18).[9] 3-azidopyridine-4-carboxylic acid (1 equivalency, 150 mg, 0.91 mmol),

1-ethyl-3-(3-dimethylaminopropyl)carbodiimide hydrochloride (2 equivalencies, 350 mg, 1.82 mmols), and N-hydroxysuccinimide (1.5 equivalencies, 155 mg, 1.35 mmols) were dissolved in 10 mL of dichloromethane and stirred at room temperature for 2 hours. The reaction mixture was transferred to a separatory funnel and the organic phase was washed with 5 mL of water (3x) and 5 mL of brine (1x). The organic layer was then dried over sodium sulfate and evaporated *in vacuo* to yield (**2**) as an orange solid in 64% yield (151 mg, 0.58 mmols). $^{1}$H NMR (500 MHz, DMSO-$d_6$): δ 8.92 (s, 1H), 8.60 (d, 1H, J = 5.0 Hz), 7.85 (d, 1H, J = 5.0 Hz), 2.90 (s, 4H). $^{13}$C NMR (500 MHz, DMSO-$d_6$): δ 169.44, 159.26, 145.87, 144.35, 135.78, 123.38, 122.37, 25.59.

*Synthesis of 2-azidobenzoic acid (3):* The synthesis of **(3)** was adapted from previous reports with minor modifications (Supplementary Figures S2, S19-20).[8] 2-aminobenzoic acid (*o*ABZ, 1 equivalency, 500 mg, 3.65 mmols) was dissolved in 10 mL of 10% HCl and stirred on ice for 10 minutes forming a partially insoluble suspension. Sodium nitrite (1.2 equivalencies, 302 mg, 4.38 mmols) was dissolved in 1.5 mL of water and slowly added to the suspension of 2-aminobenzoic acid. The resulting solution was allowed to stir at room temperature for 15 minutes. Then, sodium azide (1.2 equivalencies, 285 mg, 4.38 mmols) was dissolved in 1 mL of water and carefully added to the solution resulting in vigorous precipitation. The reaction was stirred at room temperature for 15 minutes. The precipitates were filtered, washed with water, and dried to yield **(3)** as a white solid in 33% yield (195 mg, 1.20 mmols). $^{1}$H NMR (500 MHz, DMSO-$d_6$): δ 13.14 (s, 1H), 7.77 (m, 1H), 7.60 (m, 1H), 7.36 (m, 1H), 7.26 (m, 1H,). $^{13}$C NMR (500 MHz, DMSO-$d_6$): δ 166.46, 138.68, 133.05, 131.08, 124.94, 123.97, 120.85.

*Synthesis of 2-azidobenzoic acid-N-hydroxysuccinimide (NHS) ester (4):* The synthesis of **(4)** was adapted from previous reports with minor modifications (Supplementary Figures S2, S21-22).[9] 2-azidobenzoic acid (1 equivalency, 163 mg, 1 mmol), 1-ethyl-3-(3-dimethylaminopropyl)carbodiimide hydrochloride (2 equivalencies, 385 mg, 2 mmols), and N-hydroxysuccinimide (1.5 equivalencies, 173 mg, 1.5 mmols) were dissolved in 10 mL of dichloromethane and stirred at room temperature for 2 hours. The reaction mixture was transferred to a separatory funnel and the organic phase was washed with 5 mL of water (3x) and 5 mL of brine (1x). The organic layer was then dried over sodium sulfate and evaporated *in vacuo* to yield **(4)** as a beige solid in 81% yield (210 mg, 0.81 mmols). $^{1}$H NMR (500 MHz, DMSO-$d_6$): δ 8.00 (dd, 1H, J = 8.0, 1.5 Hz), 7.83 (m, 1H), 7.59 (dd, 1H, J = 8.5, 0.5 Hz), 7.40

(m, 1H), 2.89 (s, 4H). $^{13}C$ NMR (500 MHz, DMSO-$d_6$): δ 170.27, 159.85, 140.88, 135.95, 131.72, 125.23, 121.26, 116.13, 25.56.

*Synthesis of 3-azidobenzoic acid (5):* The synthesis of **(5)** was adapted from previous reports with minor modifications (Supplementary Figure S3, S23-24).[8] 3-aminobenzoic acid (*m*ABZ, 1 equivalency, 500 mg, 3.65 mmols) was dissolved in 10 mL of 10% HCl and stirred on ice for 10 minutes to form a partially insoluble suspension. Sodium nitrite (1.2 equivalencies, 302 mg, 4.38 mmols) was dissolved in 1.5 mL of water and slowly added to the suspension of 2-aminobenzoic acid. The resulting solution was allowed to stir at room temperature for 15 minutes. Then, sodium azide (1.2 equivalencies, 285 mg, 4.38 mmols) was dissolved in 1 mL of water and carefully added to the solution resulting in vigorous precipitation. The reaction was stirred at room temperature for 15 minutes. The precipitates were filtered, washed with water, and dried to yield **(5)** as a white solid in 77% yield (460 mg, 2.82 mmols). $^{1}H$ NMR (500 MHz, DMSO-$d_6$): δ 13.23 (s, 1H), 7.74 (m, 1H), 7.54 (m, 2H), 7.36 (m, 1H). $^{13}C$ NMR (500 MHz, DMSO-$d_6$): δ 166.49, 139.92, 132.64, 130.36, 125.87, 123.48, 119.40.

*Synthesis of 3-azidobenzoate-N-hydroxysuccinimide (NHS) ester (6):* The synthesis of **(6)** was adapted from previous reports with minor modifications (Supplementary Figure S3, S25-26).[9] 3-azidobenzoic acid (1 equivalency, 163 mg, 1 mmol), 1-ethyl-3-(3-dimethylaminopropyl)carbodiimide hydrochloride (2 equivalencies, 385 mg, 2 mmols), and N-hydroxysuccinimide (1.5 equivalencies, 173 mg, 1.5 mmols) were dissolved in 10 mL of dichloromethane and stirred at room temperature for 2 hours. The reaction mixture was transferred to a separatory funnel and the organic phase was washed with 5 mL of water (3x) and 5 mL of brine (1x). The organic layer was then dried over sodium sulfate and dried *in vacuo* to yield **(6)** as a semi-beige solid in 73% yield (189 mg, 0.73 mmols). $^{1}H$ NMR (500 MHz, DMSO-$d_6$): δ 7.90 (m, 1H), 7.69 (m, 2H), 7.59 (m, 1H), 2.90 (s, 4H). $^{13}C$ NMR (500 MHz, DMSO-$d_6$): 170.21, 161.15, 140.98, 131.28, 126.35, 126.18, 126.06, 119.98, 25.56.

*Acylation of 3'-$NH_2$-$tRNA^{Phe}$* The synthesis scheme for acylated 3'-$NH_2$-$tRNA^{Phe}$ is detailed in Supplementary Figure S4. A sample of 3'-$NH_2$-$tRNA^{Phe}$ (~3,000 pmols) was diluted to 500 µL with nuclease free water and concentrated using Amicon Ultra-0.5 mL centrifugal filters with a 3 kDa molecular weight cut off according to the manufacturer's instructions. The concentrated sample was washed with 450 µL of nuclease free water (3x) to remove residual 2'-amino-2'-deoxyadenosine-5'-triphosphate. The concentration of washed 3'-$NH_2$-$tRNA^{Phe}$ was quantified *via* its absorbance at 260 nm ($A_{260}$) measured using a NanoDrop ND-1000 (Thermo Scientific) and diluted with nuclease free water to a concentration of 50 µM. Stock solutions of NHS esters **(2)**, **(4)**, and **(6)** were prepared at 50 mM in dimethyl sulfoxide. For acylation reactions, stock solution of 3'-$NH_2$-$tRNA^{Phe}$ at 50 µM in nuclease free water (20 µL, 1,000 pmols), 1 M HEPES pH 8.0 (4 µL), nuclease free water (16 µL), dimethyl sulfoxide (36 µL) and appropriate NHS ester at 50 mM in dimethyl sulfoxide (4 µL, 200 nmols) were combined. The reaction was carried out at room temperature for 16 hours. To reduce the azide-functionalized acylated $tRNA^{Phe}$, 80 µL of tris(2-carboxyethyl)phosphine (TCEP) prepared at 100 mM in 500 mM Tris pH 7.4 was added to the acylation reaction. Azide reduction was carried out at room temperature for 30 minutes. To the resulting amino-functionalized acylated $tRNA^{Phe}$, 160 µL of 600 mM sodium acetate pH 5.2 was added. Amino-functionalized acylated $tRNA^{Phe}$ was precipitated by the addition of 960 µL of ethanol and incubation at -80 °C for 1 hour. Precipitated tRNA was pelleted by centrifugation at 4 °C, 21,300 x g for 30 minutes. The pellet of amino-functionalized acylated $tRNA^{Phe}$ was suspended in 40 µL of 10 mM $NH_4OAc$, 1.7 M $(NH_4)_2SO_4$, pH 6.3 for purification via high performance liquid chromatography (HPLC).

*Purification of Acylated 3'-$NH_2$-$tRNA^{Phe}$* HPLC purification was performed on an Agilent 1260 Infinity HPLC system equipped with a UV diode array detector and an 1260 Infinity fraction collector using a 7.5 cm x 7.5 mm TSKgel™ Phenyl-5PW Column from Tosoh Bioscience (part# 807573). The mobile phase for HPLC was 10 mM $NH_4OAc$, 1.7 M $(NH_4)_2SO_4$, pH 6.3 (solvent A) and 10 mM $NH_4OAc$, pH 6.3 with 10% (v/v) methanol (solvent B) with a solvent flow rate of 0.7 mL/min. Acylated $tRNA^{Phe}$ was separated from 3'-$NH_2$-$tRNA^{Phe}$ via a linear solvent gradient from 0% to 30% solvent B over 30 minutes. Elution of tRNA species was monitored via the UV absorbance signals at 260 nm and 280 nm. Acylated $tRNA^{Phe}$ was concentrated using Amicon Ultra-0.5 mL centrifugal filters with a 3 kDa molecular weight cut off according to the manufacturer's instructions. The concentrated sample was washed with 450 µL of nuclease free water (4x). The concentration of washed 3'-amino tailed $tRNA^{Phe}$ was quantified via A260 measured using a NanoDrop ND-1000 (Thermo Scientific). Typical recovery of acylated $tRNA^{Phe}$ was approximately 200 pmols.

*Characterization of Acylated 3'-$NH_2$-$tRNA^{Phe}$* The identity of acylated 3'-NH-$tRNA^{Phe}$ was confirmed as previously described.[10] Samples were resolved on a ACQUITY UPLC BEH C18 Column (130 Å, 1.7 μm, 2.1 mm X 50 mm, Waters part # 186002350, 60 °C) using an ACQUITY UPLC I-Class PLUS (Waters part # 186015082). The mobile phases used were 8 mM triethylamine (TEA), 80 mM hexafluoroisopropanol (HFIP), 5 μM ethylenediaminetetraacetic acid (EDTA, free acid) in MilliQ water (solvent A); and 4 mM TEA, 40 mM HFIP, 5 μM EDTA (free acid) in 50% MilliQ water/50% methanol (solvent B). The method used a flow rate of 0.3 mL/min and began with Mobile Phase B at 22% that increased linearly to 40 % B over 10 min, followed by a linear gradient from 40 to 60% B for 1 min, a hold at 60% B for 1 min, a linear gradient from 60 to 22% B over 0.1 min, then a hold at 22% B for 2.9 min. The mass of the RNA was analyzed using LC-HRMS with a Waters Xevo G2-XS Tof (Waters part #186010532) in negative ion mode with the following parameters: capillary voltage: 2000 V, sampling cone: 40, source offset: 40, source temperature: 140 °C, desolvation temperature: 20 °C, cone gas flow: 10 L/h, desolvation gas flow: 800 L/h, 1 spectrum/s. Expected masses of oligonucleotide products were calculated using the AAT Bioquest RNA Molecular Weight Calculator. Deconvoluted mass spectra were obtained using the MaxEnt software (Waters Corporation).

*Cryo-EM complex preparation*

Briefly, the 70S complexes were formed by incubating 100 nM 70S ribosomes (prepared as described in Watson *et al.* 2020 [4]; gift from Fred Ward, Cate lab), 5 μM mRNA encoding Met in the P site and Phe in the A site (IDT) (Table S3), 1 μM each of fMet-NH-$tRNA^{fMet}$ and ABZ-NH-$tRNA^{Phe}$ (Table S3), and 100 μM paromomycin (Sigma) in Buffer AC (20 mM Tris pH 7.5, 100 mM $NH_4Cl$, 15 $MgCl_2$, 0.5 mM EDTA, 2 mM DTT, 2 mM spermidine, 0.05 mM spermine) in the case of the *m*ABZ complex or Grid Freezing Buffer (20 mM HEPES-KOH pH 7.5, 50 mM $NH_4Cl$, 50 mM KCl, 15 $MgCl_2$, 2 mM DTT) in the case of *o*ABZ and Apy containing complexes, for 30 min at 37 °C. The complexes were kept on ice prior to plunge freezing grids.

*Cryo-EM sample preparation*

The samples were prepared for imaging on 300 mesh R1.2/1.3 UltrAuFoil grids (Quantifoil) with an additional layer of float-transferred amorphous carbon support film. The grids were washed in chloroform prior to carbon floating. Before applying the sample, grids were glow discharged in a PELCO easiGlow at 0.37 mBar and 25 mAmp for 12 seconds. 4 μL of sample was deposited onto each grid and incubated for 1 minute. Prior to freezing, the grids were washed by successively touching three 100 μL drops of buffer containing 20 mM Tris pH 7.5, 20 mM $NH_4Cl$, 15 $MgCl_2$, 0.5 mM EDTA, 2 mM DTT, 2 mM spermidine, and 0.05 mM spermidine for *m*ABZ containing complexes, or 20 mM HEPES-KOH pH 7.5 10 mM $NH_4Cl$ 10 mM KCl, 15 mM $MgCl_2$ and 2 mM DTT for *o*ABZ and Apy containing complexes. Grids were blotted and plunge-frozen in liquid ethane with an FEI Mark IV Vitrobot using the following settings: 4°C, 100% humidity, blot force 3, and blot time 2. Grids were clipped for autoloading and stored in liquid nitrogen.

*Cryo-EM data collection*

Cryo-EM data collection parameters are summarized in Supplementary Tables S1-S3. For the *o*ABZ and Apy complexes, dose fractionated movies were collected on a Titan Krios G3i microscope at an accelerating voltage of 300 kV and with a BIO Quantum energy filter. In the case of *m*ABZ, a Titan Krios G2 microscope with a BIO Quantum energy filter (300 kV) was used. Movies were recorded on a GATAN K3 direct electron detector operated in CDS mode. A total dose of 40 $e^-/Å^2$ was split over 40 frames per movie. The physical pixel size was set to 0.81 Å and the super-resolution pixel size to 0.405 Å in the case of the *o*ABZ and Apy complexes. The pixel size for the *m*ABZ dataset was 0.71 Å with the super-resolution pixel size being 0.355 Å. Data collection was automated with SerialEM, which was also used for astigmatism correction by CTF and coma-free alignment by CTF. For the *o*ABZ and *m*ABZ complexes, stage shift was used to move between center holes of a 3×3 hole template and image shift to collect on the surrounding 8 holes. One movie per hole was collected for the *m*ABZ complex, and three movies per hole were collected for the *o*ABZ using image shift. In the case of the Apy complex, stage shift was used to move between the center holes of a 7×7 hole template, and image shift was used to collect on the surrounding 48 holes. Image shift was also used to collect two movies per hole in this case. The defocus ramp was set to range between -0.5 and -1.5 μm.

*Image Processing*

Data processing was performed using RELION[11,12] and cryoSPARC. Initial processing for *m*ABZ was done using RELION 3.152,[11] with later steps being completed in RELION 4.0.0[12]. *o*ABZ and Apy were processed primarily in RELION 4.0.0, with the exception of initial 3D classification in cryoSPARC[13]. MotionCor2[14] was used to motion correct the raw movies (binned to the physical pixel size estimated to be 0.81 Å during initial steps). CtfFind4[15] was used to estimate CTF parameters, and micrographs with poorly fitting CTF estimates were omitted after visual inspection. Particles were picked using the Laplacian-of-Gaussian autopicking method in RELION. Particles were then extracted at one-eighth binning of the full box size, and subjected to three rounds of 2D refinement to select 70S particles. The selected particles were then re-extracted after scaling to one-fourth of the full size and imported to cryoSPARC, where 3D classification was performed using the "Heterogeneous Refinement" job using a 70S reference volume generated from the coordinates of PDB entry 1VY4[1] using EMAN2[16]. Classes corresponding to clean 70S volumes were selected and subjected to 50S-focused refinement, and exported back to RELION. To separate classes with rotations of the 30S subunit relative to the 50S subunit, 3D classification with no alignment was performed.[17] For the *o*ABZ structure, this step yielded a single 70S class, with the remaining classes being junk classes. For the Apy and *m*ABZ structures, two classes with subtle rotations of the 30S subunit relative to the 50S subunit were observed. However, both classes in each case contained density for A-site tRNA, and were pooled. Following this, masked classification for particles containing A-site tRNA was performed (with no alignments) and particles containing strong density for A-site tRNA were selected and re-extracted at full size. The particles were refined using the 3D auto-refine job in RELION, and then subjected to CTF Refinement[18], Bayesian Polishing[19], another round of CTF Refinement, and finally 50S and 30S focused refinements to improve global resolution. The global resolutions of the final 50S focused refinement maps were 2.1 Å for Apy, 1.9 Å for *o*ABZ and 2.3 Å for *m*ABZ (Supplementary Figures S6-S8).

*Pixel Size Calibration*

The pixel size was calibrated in UCSF ChimeraX[20] using the "Fit to Map" function and the high resolution maps against the coordinates from the X-ray crystal structure PDB 4YBB[21]. The best cross-correlation value was obtained at pixel size 0.8240 Å for Apy, 0.8232 Å for *o*ABZ and 0.7125 Å for *m*ABZ. Half-maps for the refined volumes were rescaled to the calibrated pixel size in ChimeraX and post-processing was performed with these half-maps in RELION to obtain the FSC curves and final resolution estimates.

*Modeling*

PDB entry 8EBB containing Met-tRNA$^{fMet}$ in both the P site and the A site was used as the starting model. The current structures contained tRNA$^{Phe}$ in the A site, and these coordinates were obtained from PDB 1VY4[1]. The aminobenzoic acid derivatives were modeled in Avogadro[22], and restraints for these monomers were generated using the Prodrug Server[23]. Real space refinement of the coordinates was performed in PHENIX[24] and further adjustments to the model were done manually in Coot[25], mainly in the vicinity of the PTC. Further additions to the model included $Mg^{2+}$ and $K^{+}$ ions and water molecules. The linkage between A- and P-site tRNAs and monomers is modeled as an amide linkage to reflect experimental conditions for structure determination. The map-vs.-model FSC was calculated in PHENIX[24].

*pKa calculations*

All three compounds were prepared as their ammonium cations and analyzed using the Schrödinger Maestro Suite (version 2022-4). pKa values were determined using Jaguar v11.8 following an initial conformational search with Macromodel (v13.8, OPLS4 force field).

**Supplementary Tables**

**Table S1:** Data collection and processing for WT *E. coli* ribosome complexes with aminobenzoic acid monomers

| Complex | Apy | *o*ABZ | *m*ABZ |
|---|---|---|---|
| Magnification | 103,216 | 103,316 | 104,632 |
| Voltage (kV) | 300 | 300 | 300 |
| Electron Exposure ($e^-$/Å$^2$) | 40 | 40 | 40 |
| Defocus Range (µm) | -0.5/-1.5 | -0.5/-1.5 | -0.5/-1.5 |
| Pixel Size (Å) | 0.8240 | 0.8232 | 0.7125 |
| Symmetry Imposed | C1 | C1 | C1 |
| Initial Particle Images | 2,104,199 | 1,593,031 | 646,976 |
| Final Particle Images | 288,552 | 339,543 | 144,049 |
| Map Resolution (Å) | 2.08 | 1.89 | 2.3 |
| FSC Threshold | 0.143 | 0.143 | 0.143 |

**Table S2**: Model Refinement Statistics for WT *E. coli* ribosome complexes with aminobenzoic acid monomers.

| Model component | Apy | *o*ABZ | *m*ABZ |
|---|---|---|---|
| Model resolution (Å) | 2.1 | 2.0 | 2.3 |
| FSC threshold | 0.5 | 0.5 | 0.5 |
| Map sharpening *B* factor (Å$^2$) | -38.8702 | -42.7031 | -41.5798 |
| Model composition | | | |
| Non-hydrogen atoms | 145457 | 145449 | 145632 |
| $Mg^{2+}$ ions | 320 | 320 | 320 |
| Waters | 3491 | 3492 | 3491 |
| Mean *B* Factors (Å$^2$) | | | |
| RNA | 31.72 | 21.56 | 42.07 |
| Protein | 23.54 | 13.30 | 24.94 |
| Waters | 6.67 | 4.38 | 7.70 |
| Other | 16.23 | 9.26 | 14.63 |
| R.m.s. deviations from ideal values | | | |
| Bond (Å) | 0.006 | 0.008 | 0.006 |

| | | | |
|---|---|---|---|
| Angle (°) | 0.926 | 0.904 | 0.980 |
| Molprobity score | 2.41 | 2.42 | 2.68 |
| Clash Score | 8.83 | 9.36 | 11.36 |
| Rotamer outliers (%) | 6.50 | 6.72 | 8.42 |
| Ramachandran plot | | | |
| Favored (%) | 95.36 | 95.78 | 93.85 |
| Allowed (%) | 4.46 | 4.13 | 5.99 |
| Outliers (%) | 0.18 | 0.09 | 0.15 |
| RNA validation | | | |
| Angles outliers (%) | 0.011 | 0.008 | 0.013 |
| Sugar pucker outliers (%) | 14.6 | 14.5 | 16.8 |
| Average suiteness | 0.561 | 0.564 | 0.530 |

**Table S3:** Sequences of RNA oligomers used in this study

| RNA name | Sequence | Notes |
|---|---|---|
| $tRNA^{fMet}$ | 5'-GGCGGGGUGGAGCAGCCUGGUAGCUCGUCGGGCUCAUAACCCGAAGGUCGUCGGUUCAAAUCCGGCCCCCGCAACCA-3' | Anticodon underlined |
| $tRNA^{Phe}$ | 5'-GCCCGGAUAGCUCAGUCGGUAGAGCAGGGGAUUGAAAAUCCCCGUGUCCUUGGUUCGAUUCCGAGUCCGGGCACCA-3' | Anticodon underlined |
| mRNA | 5'- GUAUAA**GGAGG**UAAAAAUGUUCUAACUA-3' | Shine-Dalgarno sequence in bold, Met and Phe codons underlined |

**Table S4:** Sequences of DNA primers used in this study. mG represents 2'-O-methylguanosine modification

| Primer name | Sequence |
|---|---|
| fMet_C1G_temp_Fw | 5'-AATTCCTGCAGTAATACGACTCACTATAGGCGGGGTGGAGCAGCCTGGTAGCTCGTCGGGCTCATA-3' |

| | |
|---|---|
| fMet_C1G_temp_Rv(-A) | 5'-GGTTGCGGGGGCCGGATTTGAACCGACGACCTTCGGG TTATGAGCCCGACGAGCTA-3' |
| fMet_C1G_amp_Fw | 5'-AATTCCTGCAGTAATACGACTCAC-3' |
| fMet_C1G_amp_Rv(-A) | 5'-GmGTTGCGGGGGCC-3' |
| Phe_temp_Fw | 5'-AATTCCTGCAGTAATACGACTCACTATAGCCCGGATAG CTCAGTCGGTAGAGCAGGGGAT-3' |
| Phe_temp_Rv(-A) | 5'-GGTGCCCGGACTCGGAATCGAACCAAGGACACGGGG ATTTTCAATCCCCTGCTCTACCG-3' |
| Phe_amp_Fw | 5'-AATTCCTGCAGTAATACGACTCA-3' |
| Phe_amp_Rv (-A) | 5'-GmGTGCCCGGACTCG-3' |

**Supplementary Figures**

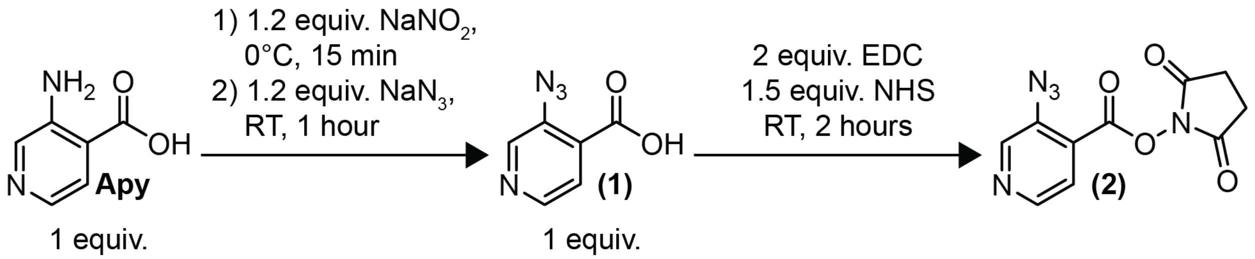


**Supplementary Figure S1.** Synthesis of 3-azidopyridine-4-carboxylic acid **(1)** and 3-azidopyridine-4-carboxylic acid-N-hydroxysuccinimide (NHS) ester **(2)**.

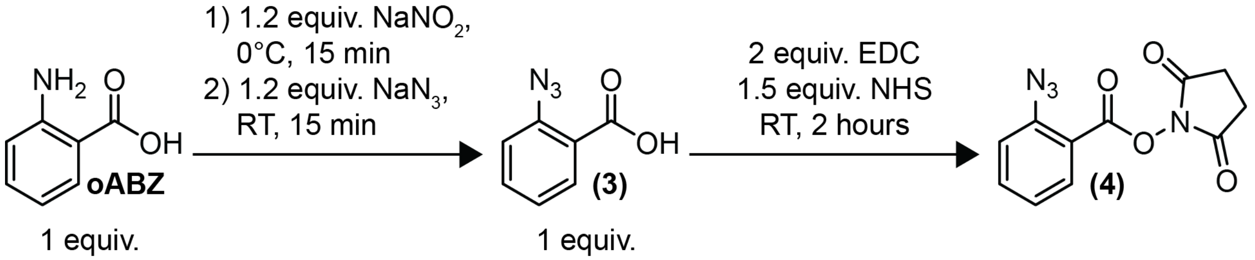


**Supplementary Figure S2.** Synthesis of 2-azidobenzoic acid **(3)** and 2-azidobenzoic acid-N-hydroxysuccinimide (NHS) ester **(4)**.

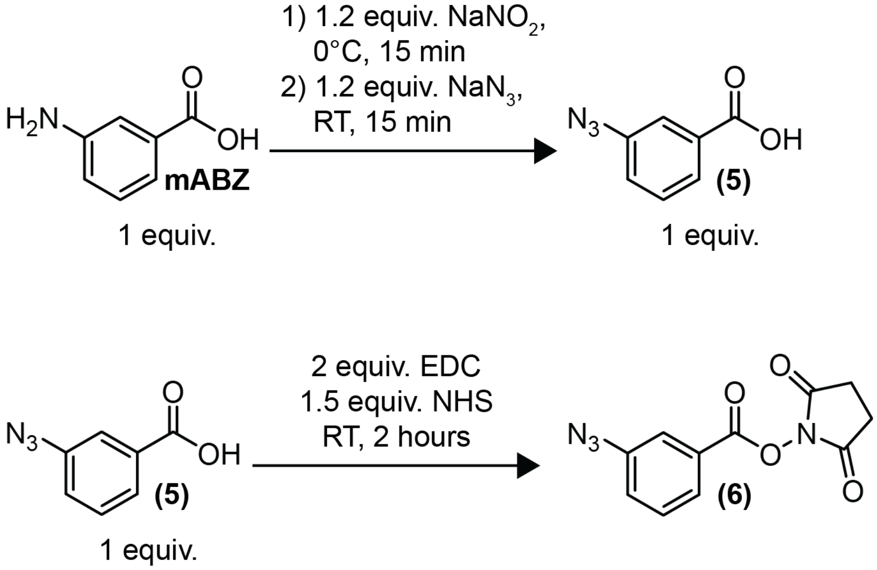


**Supplementary Figure S3.** Synthesis of 3-azidobenzoic acid **(5)** and 3-azidobenzoic acid-N-hydroxysuccinimide (NHS) ester **(6)**.

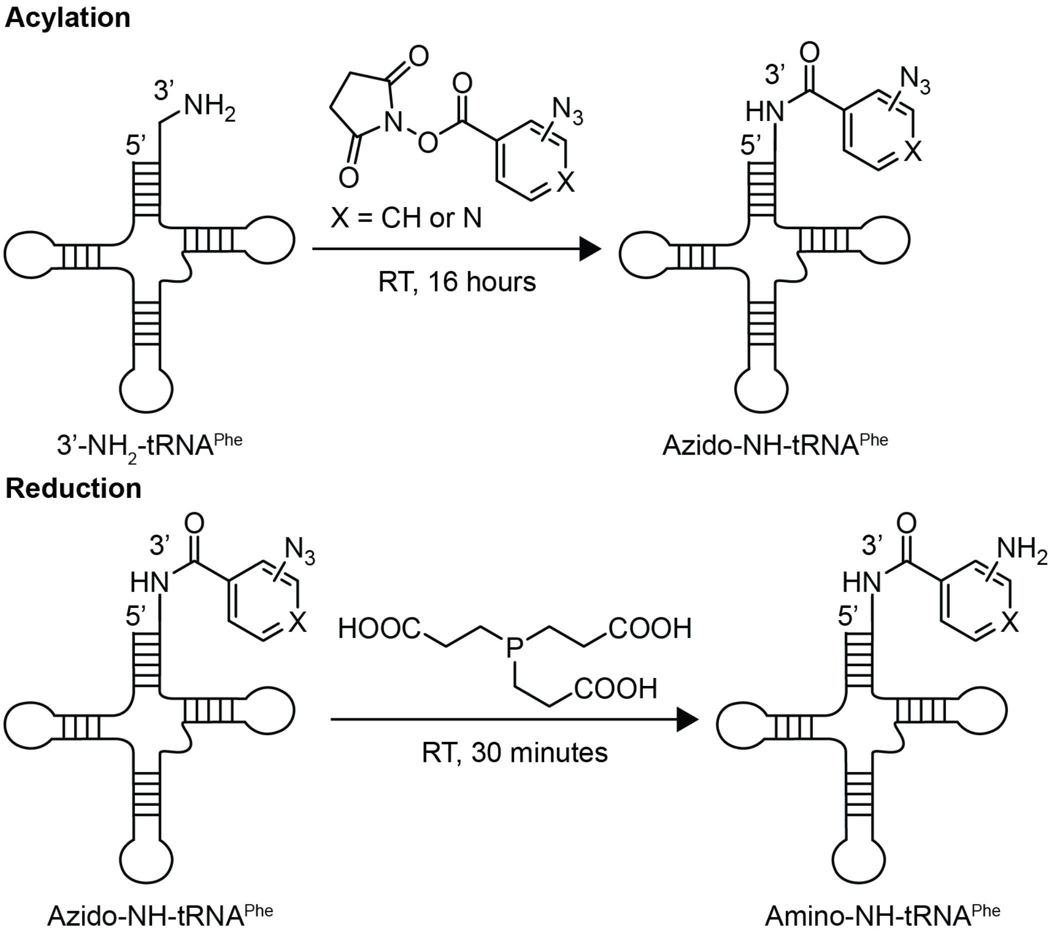


**Supplementary Figure S4.** Synthesis of acylated 3'-$NH_2$-$tRNA^{Phe}$.

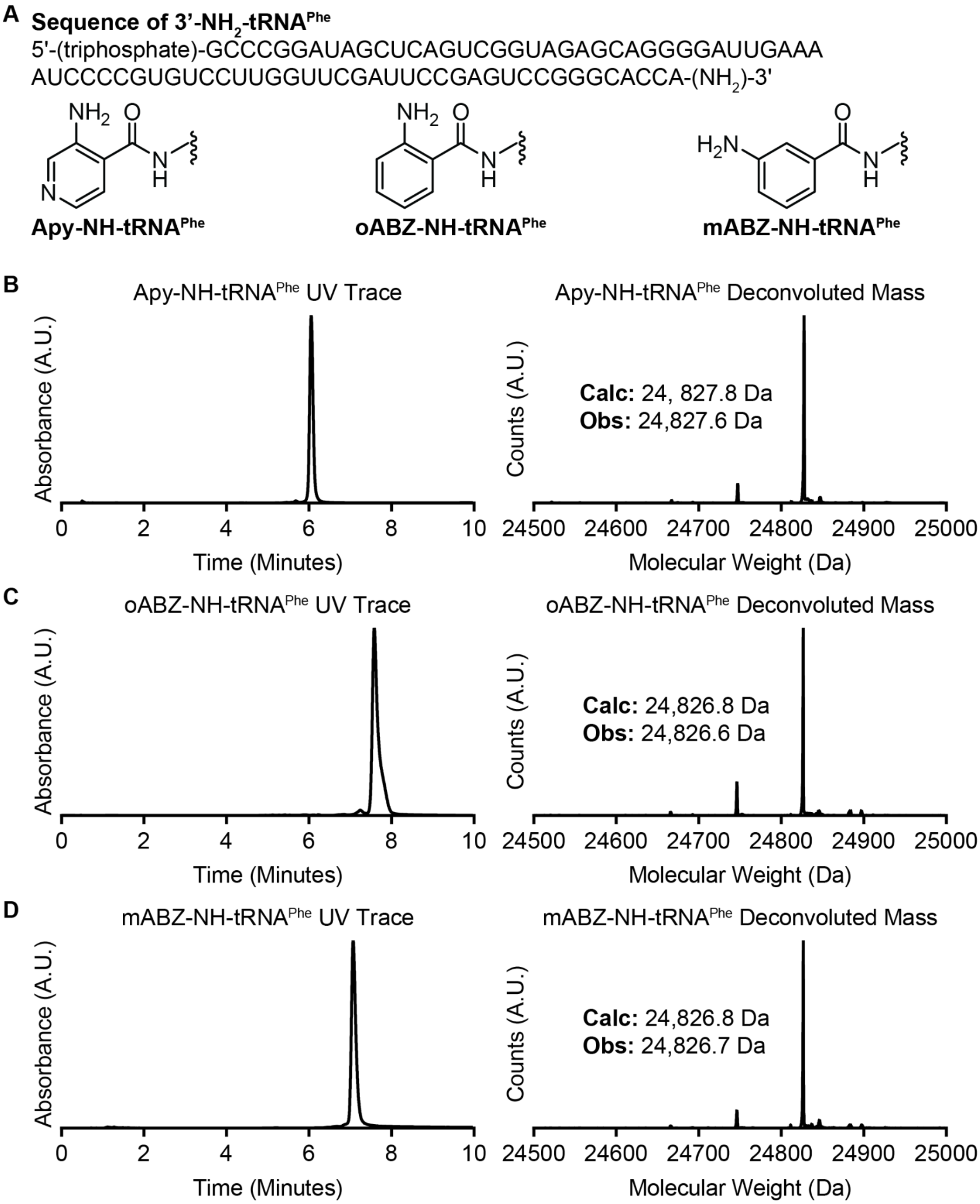


**Supplementary Figure S5.** Characterization of acylated 3'-$NH_2$-$tRNA^{Phe}$ variants after purification *via* HPLC. (A) RNA sequence and 3' modifications of acylated 3'-$NH_2$-$tRNA^{Phe}$ used in this study. (B) UV and mass spectrometry characterization of Apy-NH-$tRNA^{Phe}$. (C) UV and mass spectrometry characterization of *o*ABZ-NH-tRNA$^{Phe}$. (D) UV and mass spectrometry characterization of *m*ABZ-NH-tRNA$^{Phe}$.

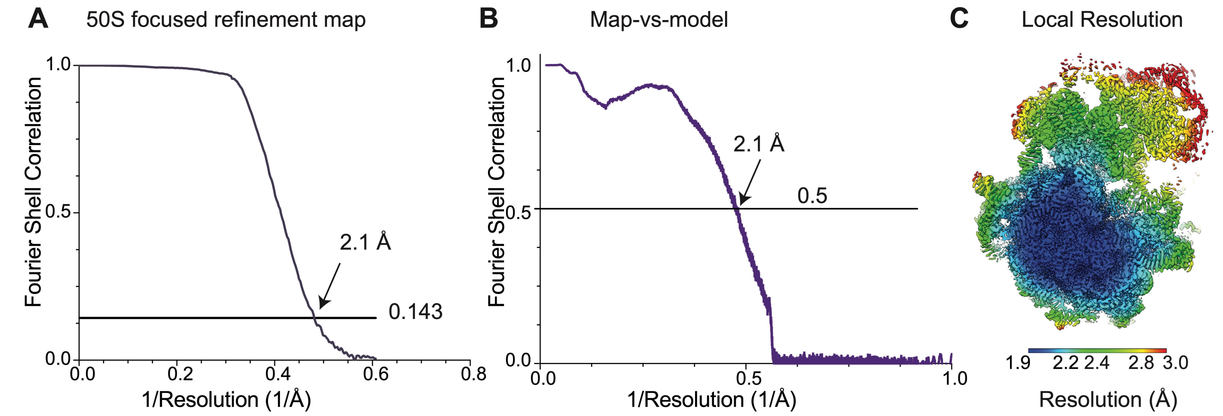


**Supplementary Figure S6**: Resolution of the 50S subunit of the fMet-Apy complex. (A) FSC curve of the 50S subunit focused map (blue) showing the global resolution of the map is 2.1 Å at the gold-standard cutoff value of 0.143. (B) Map-vs-model resolution is at 2.2 Å at FSC cutoff value of 0.5. (C) 50S focused map color coded by local resolution values.

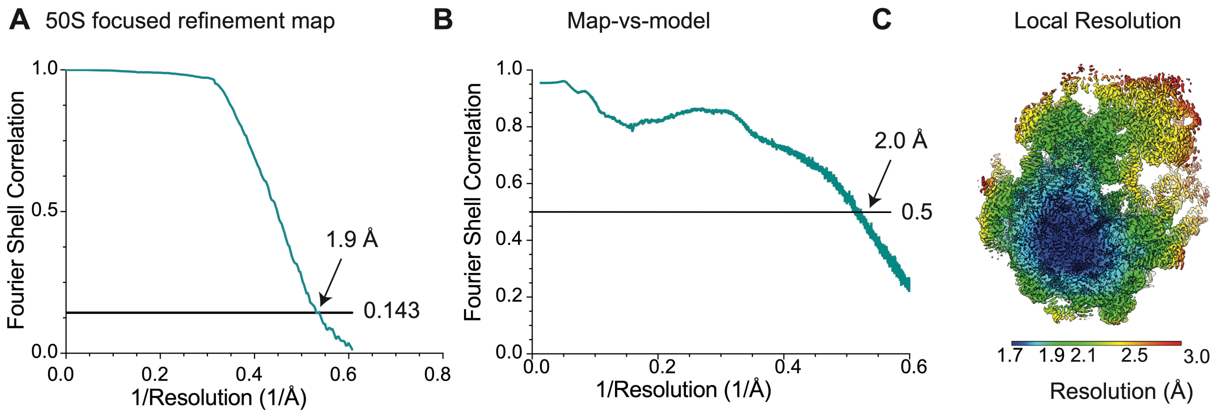


**Supplementary Figure S7**: Resolution of the 50S subunit of the fMet-*o*ABZ complex. (A) FSC curve of the 50S subunit focused map (blue) showing the global resolution of the map is 1.9 Å at the gold-standard cutoff value of 0.143. (B) Map-vs-model resolution is at 2.3 Å at FSC cutoff value of 0.5. (C) 50S focused map color coded by local resolution values.

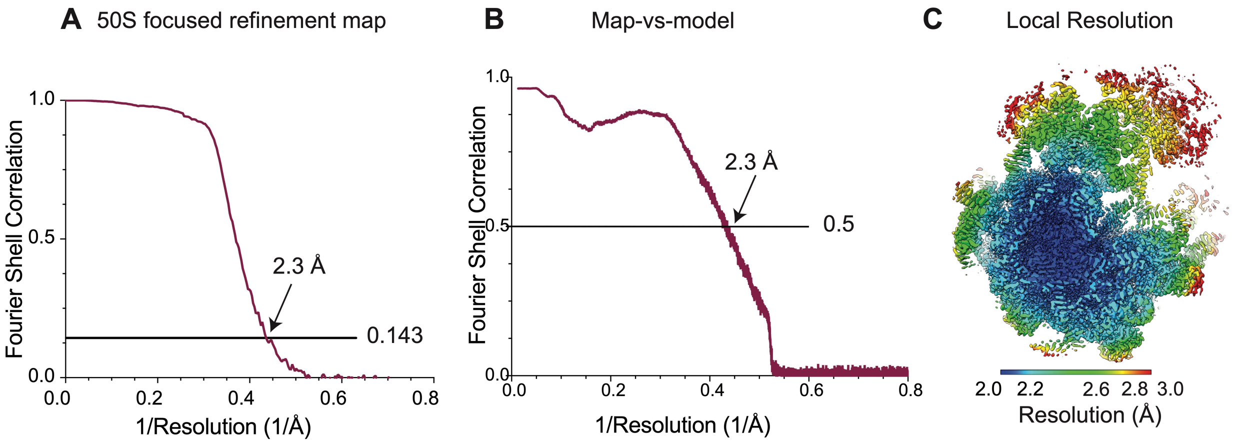


**Supplementary Figure S8**: Resolution of the 50S subunit of the fMet-*m*ABZ complex. (A) FSC curve of the 50S subunit focused map (blue) showing the global resolution of the map is 2.3 Å at the gold-standard cutoff value of 0.143. (B) Map-vs-model resolution is at 2.3 Å at FSC cutoff value of 0.5. (C) 50S focused map color coded by local resolution values.

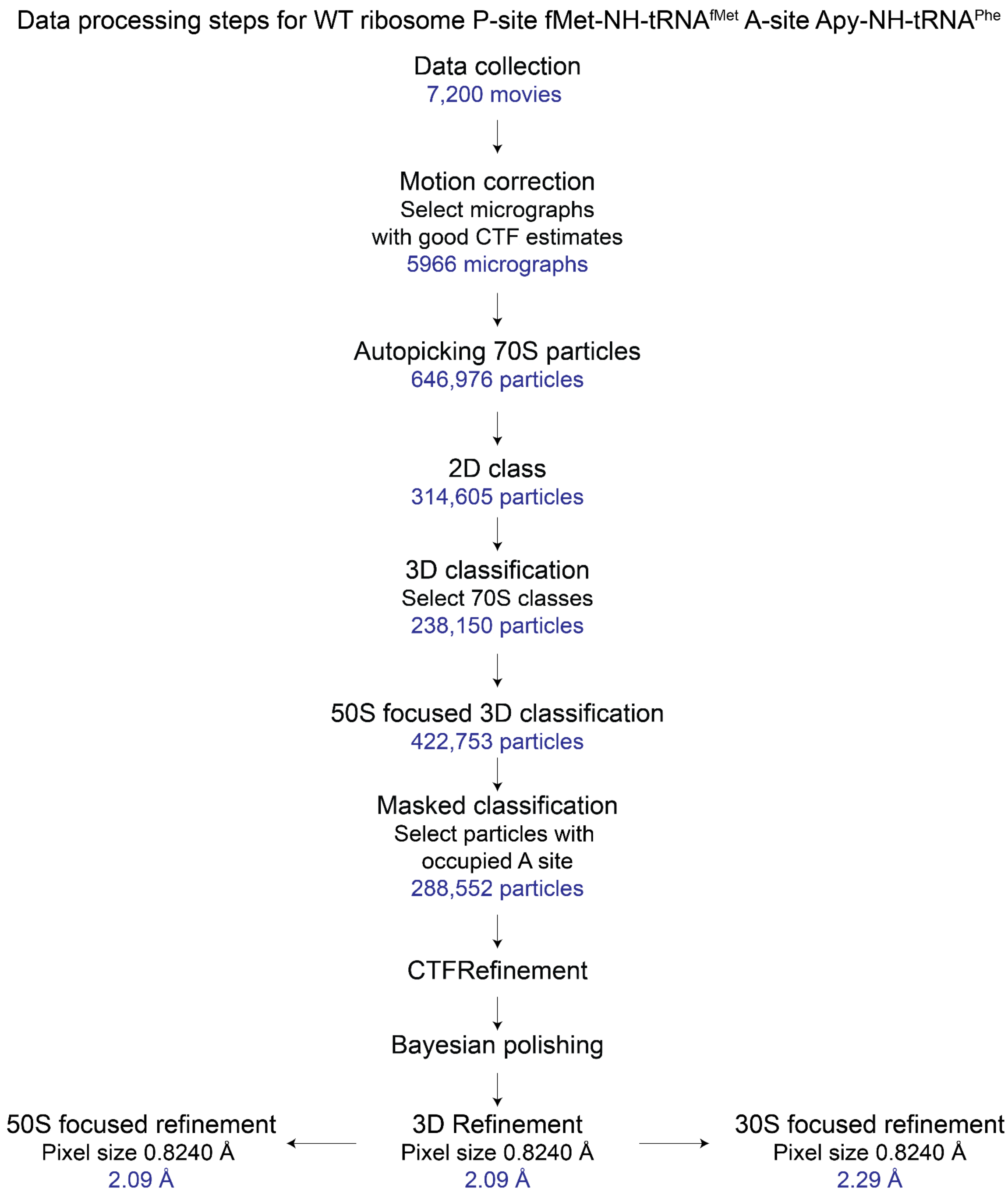


**Supplementary Figure S9**: Cryo-EM data processing workflow for the Apy containing complex. Number of movies, micrographs or particles, or resolution of the obtained volumes are indicated at each step.

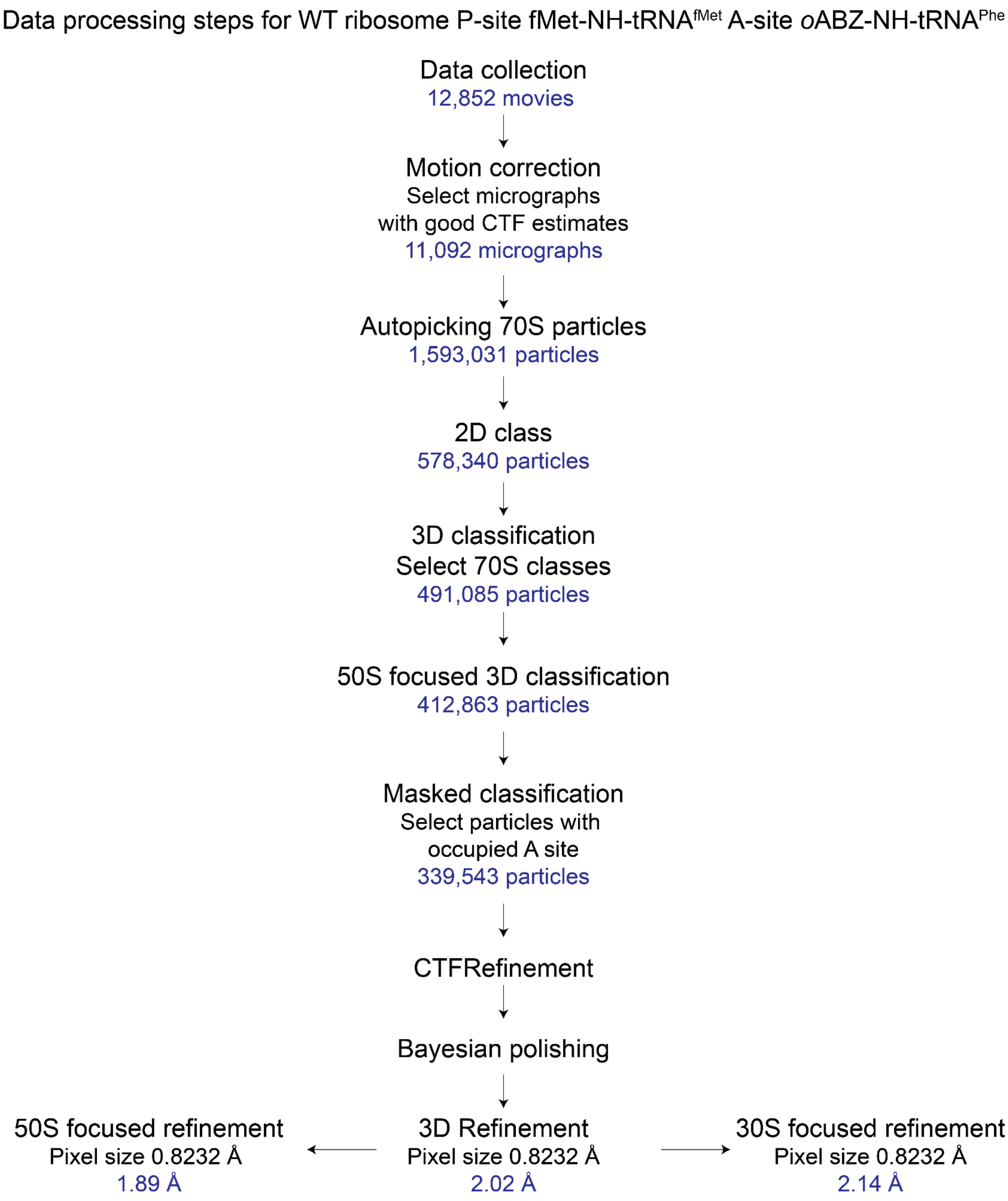


**Supplementary Figure S10**: Cryo-EM data processing workflow for the *o*ABZ containing complex. Number of movies, micrographs or particles, or resolution of the obtained volumes are indicated at each step.

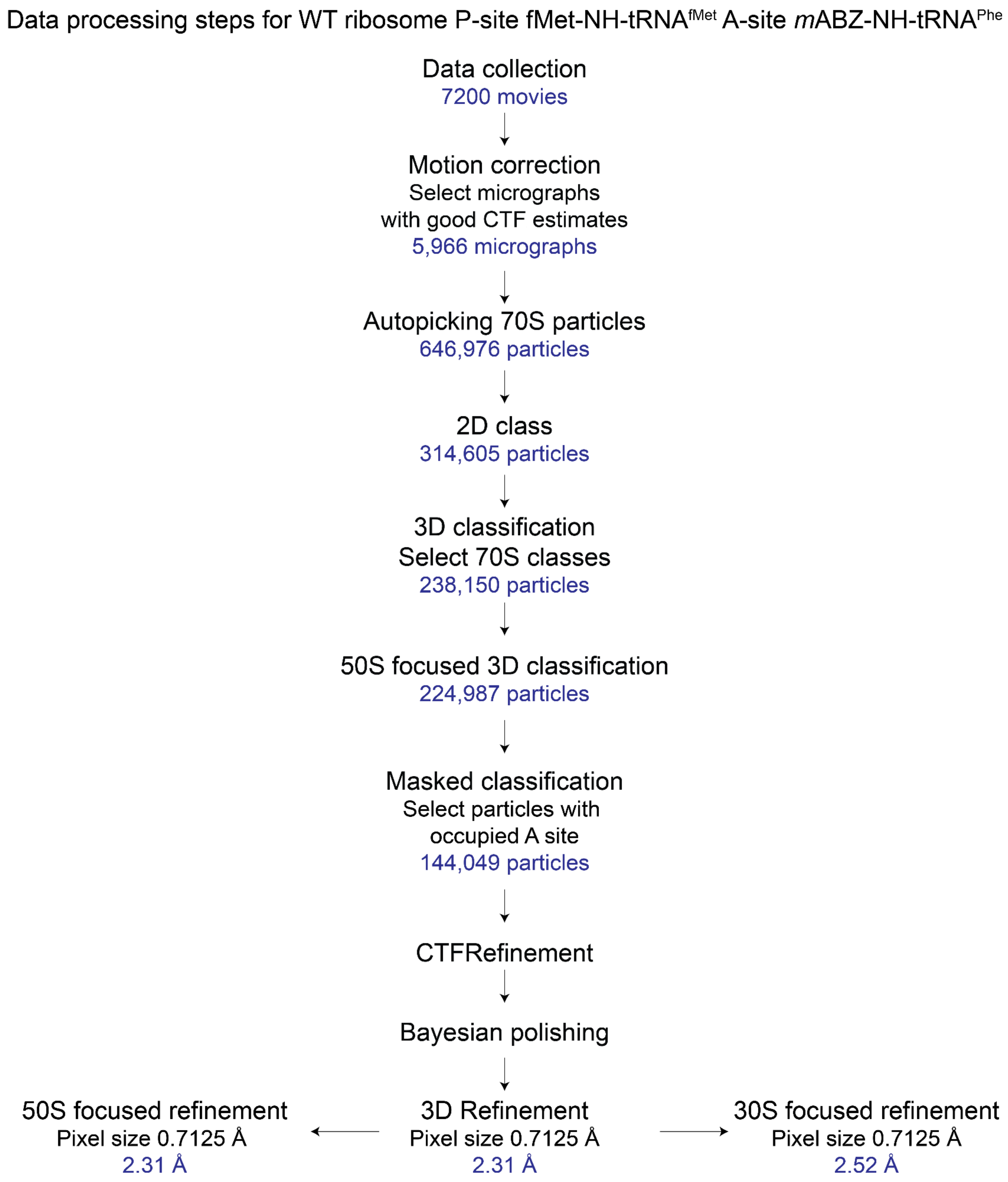


**Supplementary Figure S11**: Cryo-EM data processing workflow for the *m*ABZ containing complex. Number of movies, micrographs or particles, or resolution of the obtained volumes are indicated at each step.

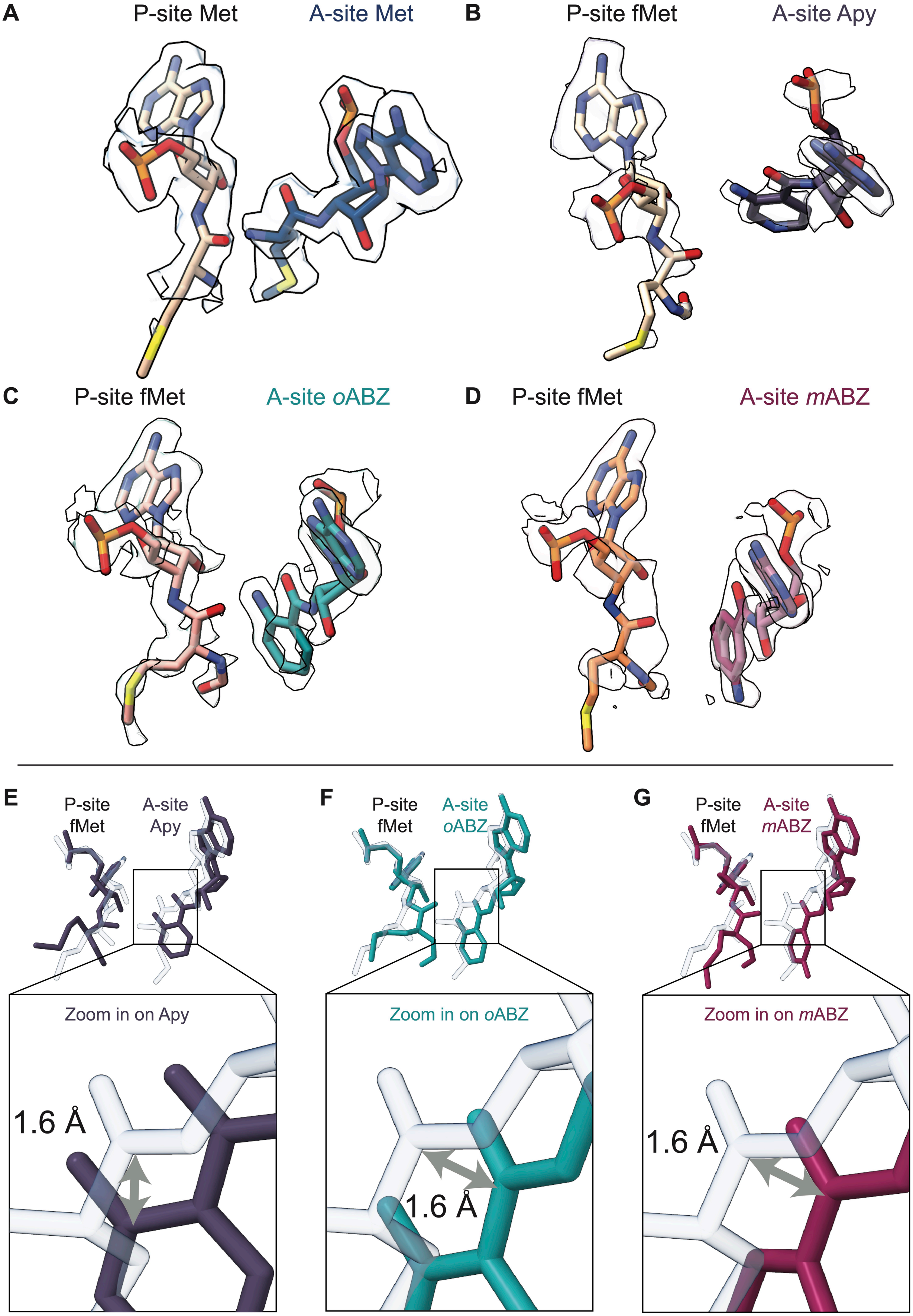
A
P-site Met
A-site Met
B
P-site fMet
A-site Apy
C
P-site fMet
A-site oABZ
D
P-site fMet
A-site mABZ
E
P-site fMet
A-site Apy
Zoom in on Apy
1.6 Å
F
P-site fMet
A-site oABZ
Zoom in on oABZ
1.6 Å
G
P-site fMet
A-site mABZ
Zoom in on mABZ
1.6 Å

**Supplementary Figure S12**: Relative positioning of the reactive groups of P- and A-site monomers showing cryo-EM density of the P- and A-site monomers. (A) P-site Met and A-site Met (B) P-site fMet and A-site Apy (C) P-site fMet and A-site *o*ABZ (D) P-site fMet and A-site *m*ABZ. The two conformations of *m*ABZ are shown in maroon and pink atomistic representation. Zoom in view showing that accommodation of the aminobenzoic acid derivatives (E) Apy, (F) *o*ABZ and (G) *m*ABZ within the A site results in a shift of the A-site carbonyl group by ~ 1.6 Å.

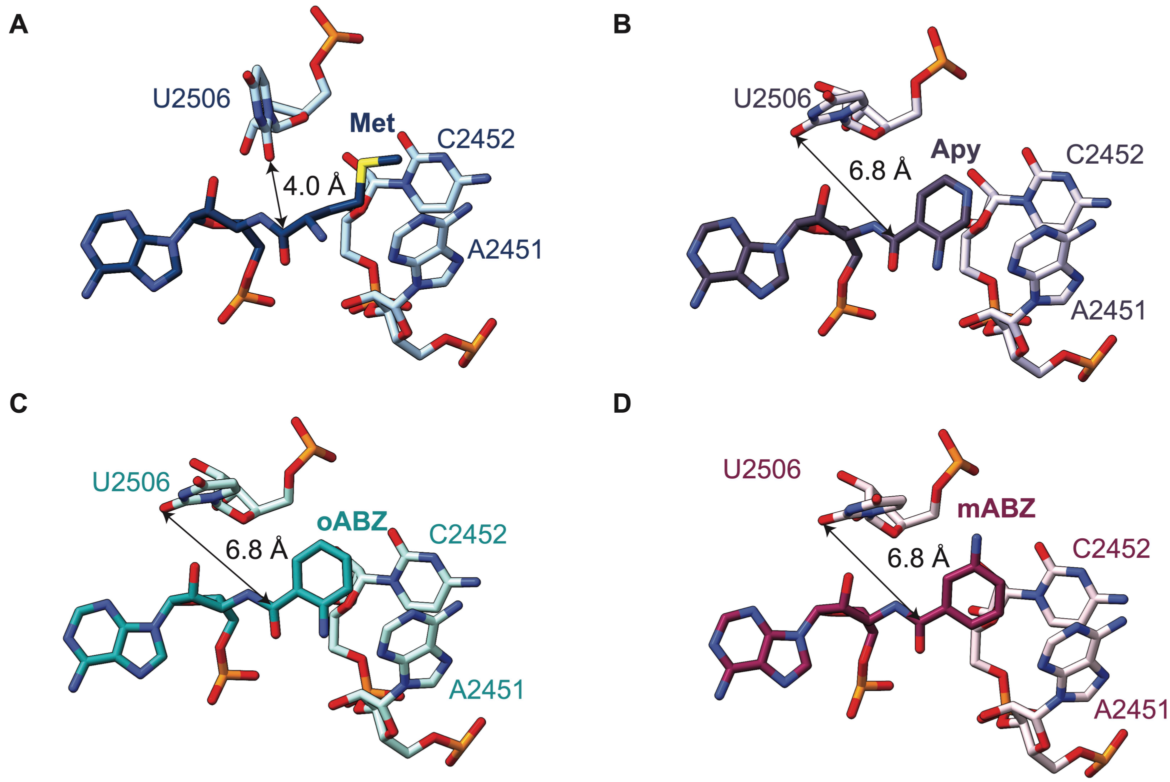


**Supplementary Figure S13**: Repositioning of nucleotide U2506 in the complexes containing aminobenzoic acid derivatives. (A) In the canonical positioning of U2506 with an A-site methionine, the O2 is positioned 4 Å away from the carbonyl carbon (PDB ID 8EBB).[5] When aminobenzoic acid derivatives are in the A site, the distance between these atoms increases to 6.8 Å as shown in the case of (B) Apy, (C) *o*ABZ, and (D) *m*ABZ.

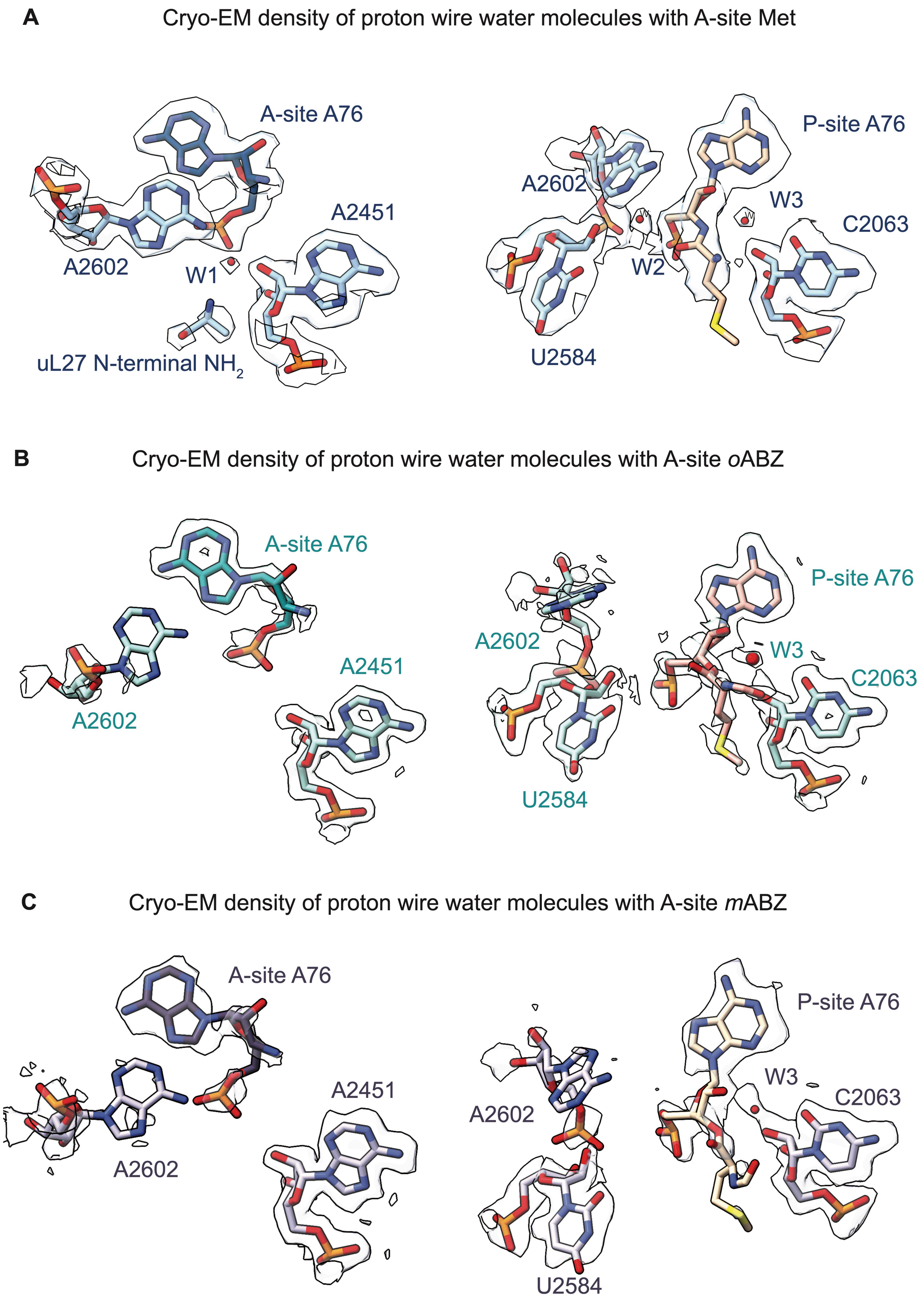

A
Cryo-EM density of proton wire water molecules with A-site Met
A-site A76
A2451
A2602
W1
uL27 N-terminal NH2
P-site A76
A2602
W3
C2063
W2
U2584
B
Cryo-EM density of proton wire water molecules with A-site oABZ
A-site A76
A2451
A2602
P-site A76
A2602
W3
C2063
U2584
C
Cryo-EM density of proton wire water molecules with A-site mABZ
A-site A76
A2451
A2602
P-site A76
W3
C2063
A2602
U2584

**Supplementary Figure S14**: Binding of aminobenzoic acid monomers in the A-site disrupts the proton wire proposed to support proton transfers during amide bond formation. (A) Position of the three water molecules associated with the proton wire (W1, W2, and W3)[1] and observed in the ribosome-tRNA complex with an A-site methionine (PDB ID 8EBB). In the complexes with a *o*ABZ (B) or Apy (C) monomer in the A site, nucleotide A2602 and the N-terminal tail of protein uL27 are disordered and no cryo-EM density for W1 or W2 is observed. Density corresponding to W3 appears in between the 2'-OH groups of nucleotides C2063 and A76 of the P-site tRNA. Maps were supersampled for smoothness.

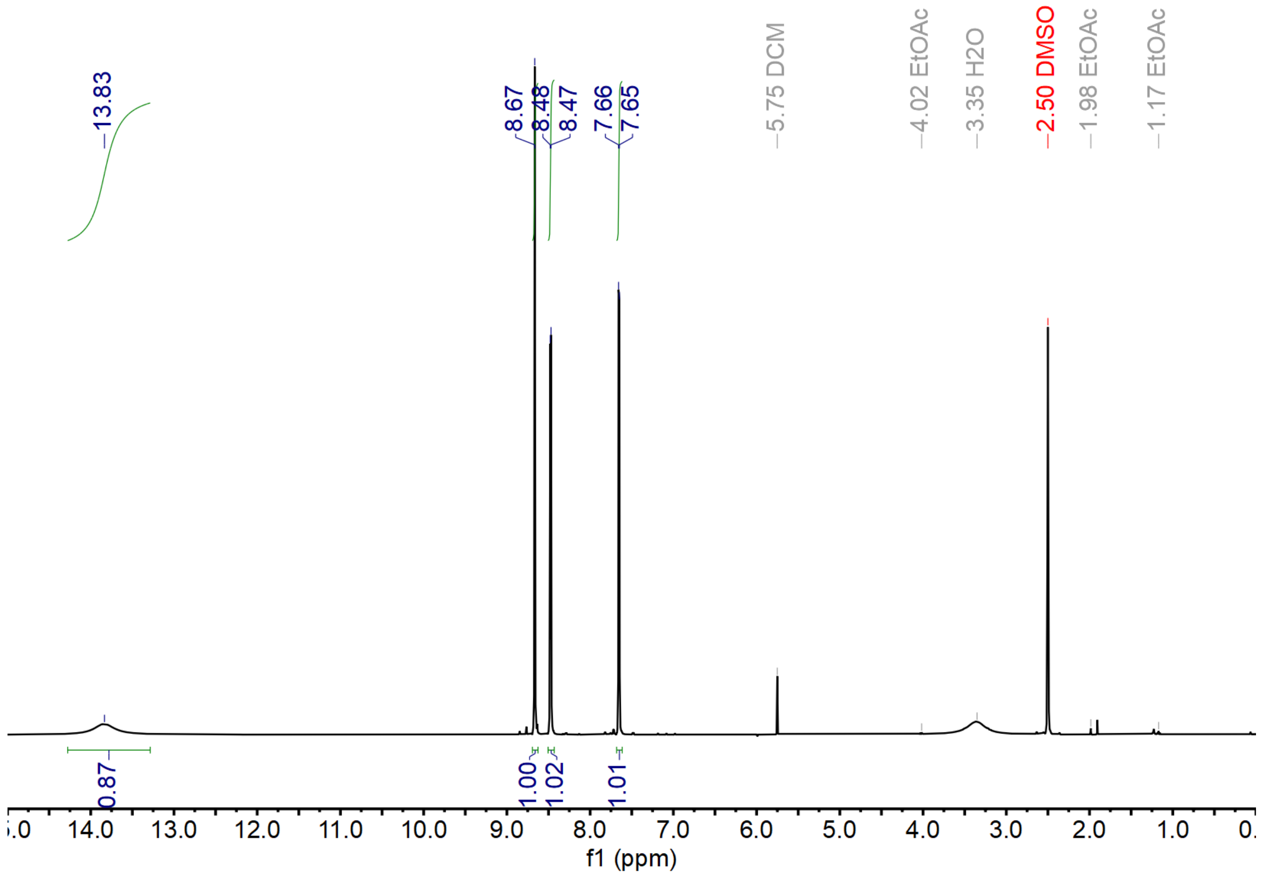


**Supplementary Figure 15.** $^{1}$H NMR (500 MHz, DMSO-$d_6$) characterization of compound **(1)**.

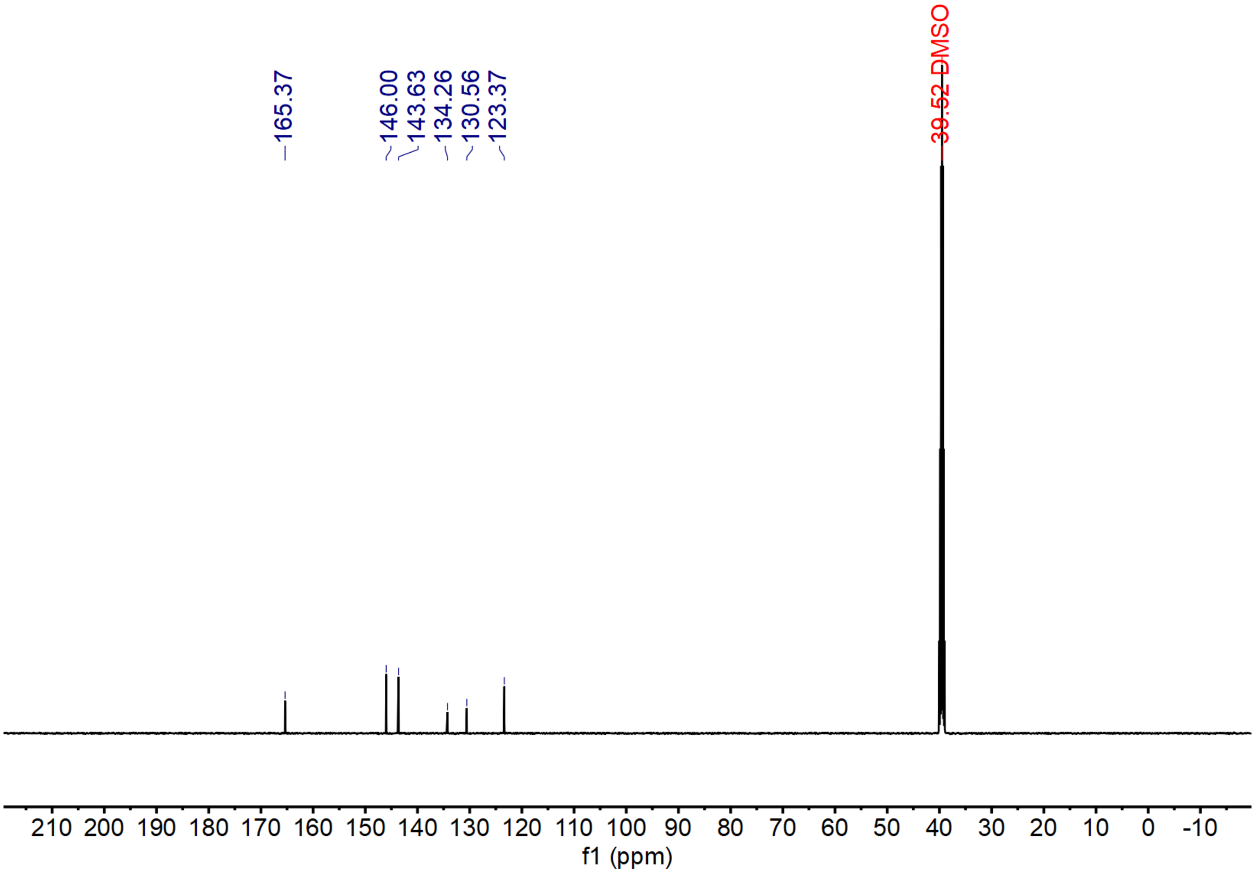


**Supplementary Figure 16.** $^{13}C$ NMR (500 MHz, DMSO-$d_6$) characterization of compound **(1)**.

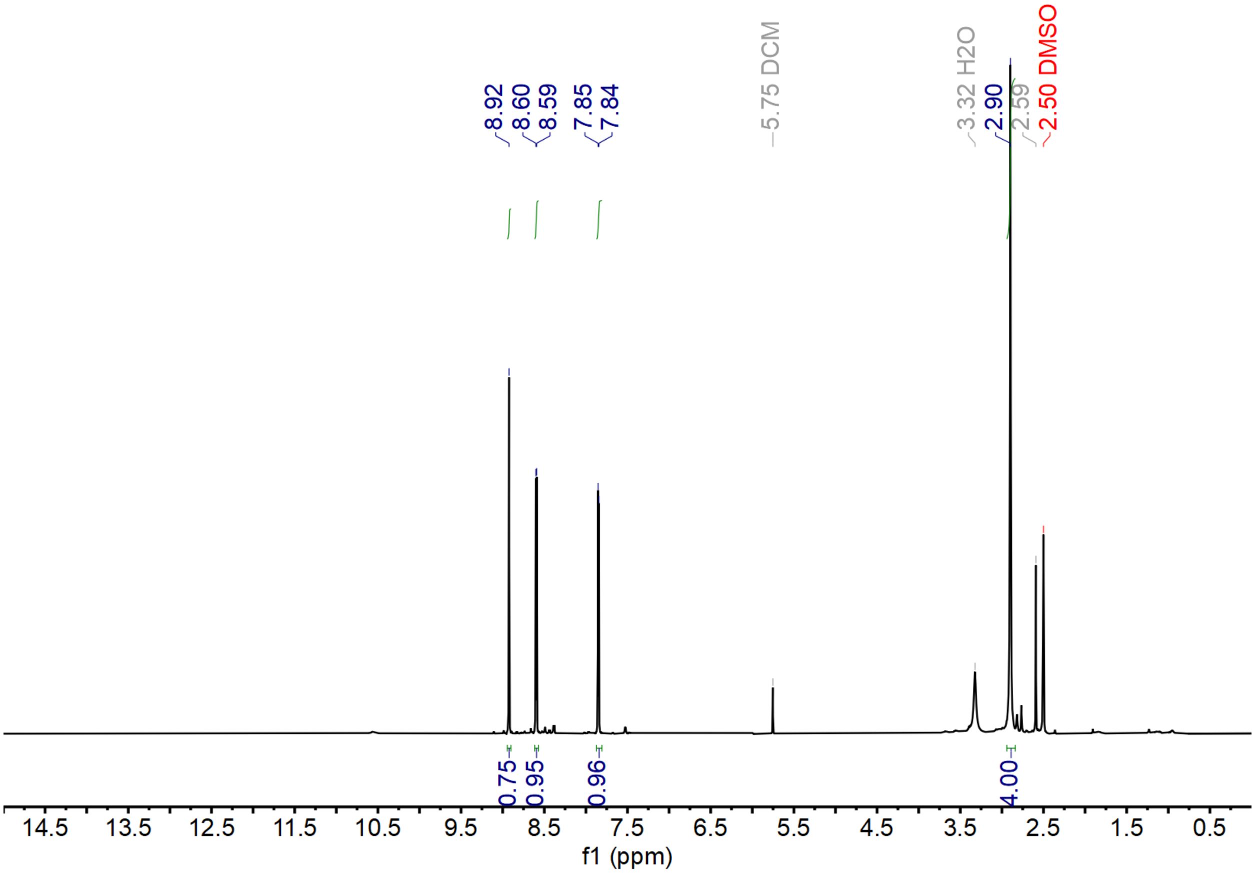


**Supplementary Figure 17.** $^1$H NMR (500 MHz, DMSO-$d_6$) characterization of compound **(2)**.

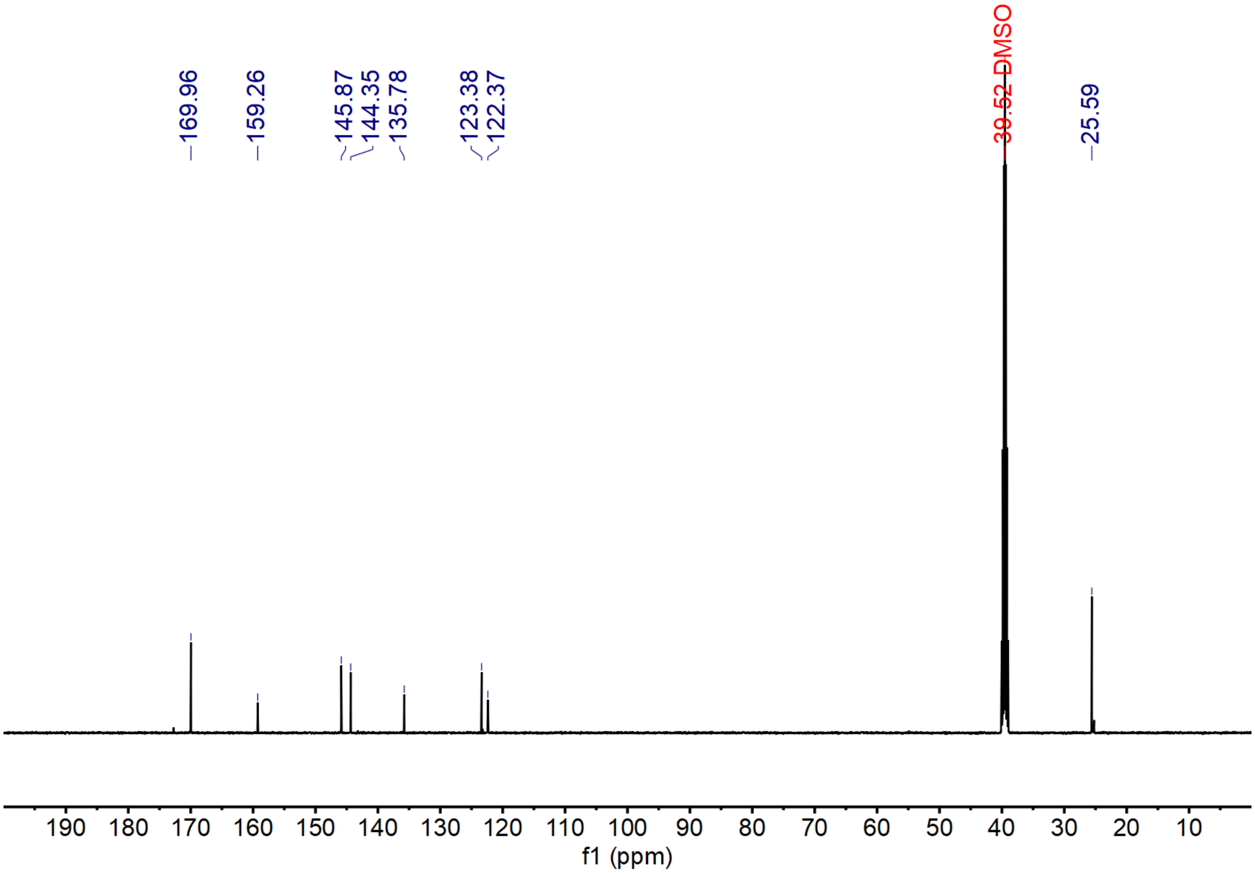


**Supplementary Figure 18.** $^{13}C$ NMR (500 MHz, DMSO-$d_6$) characterization of compound **(2)**.

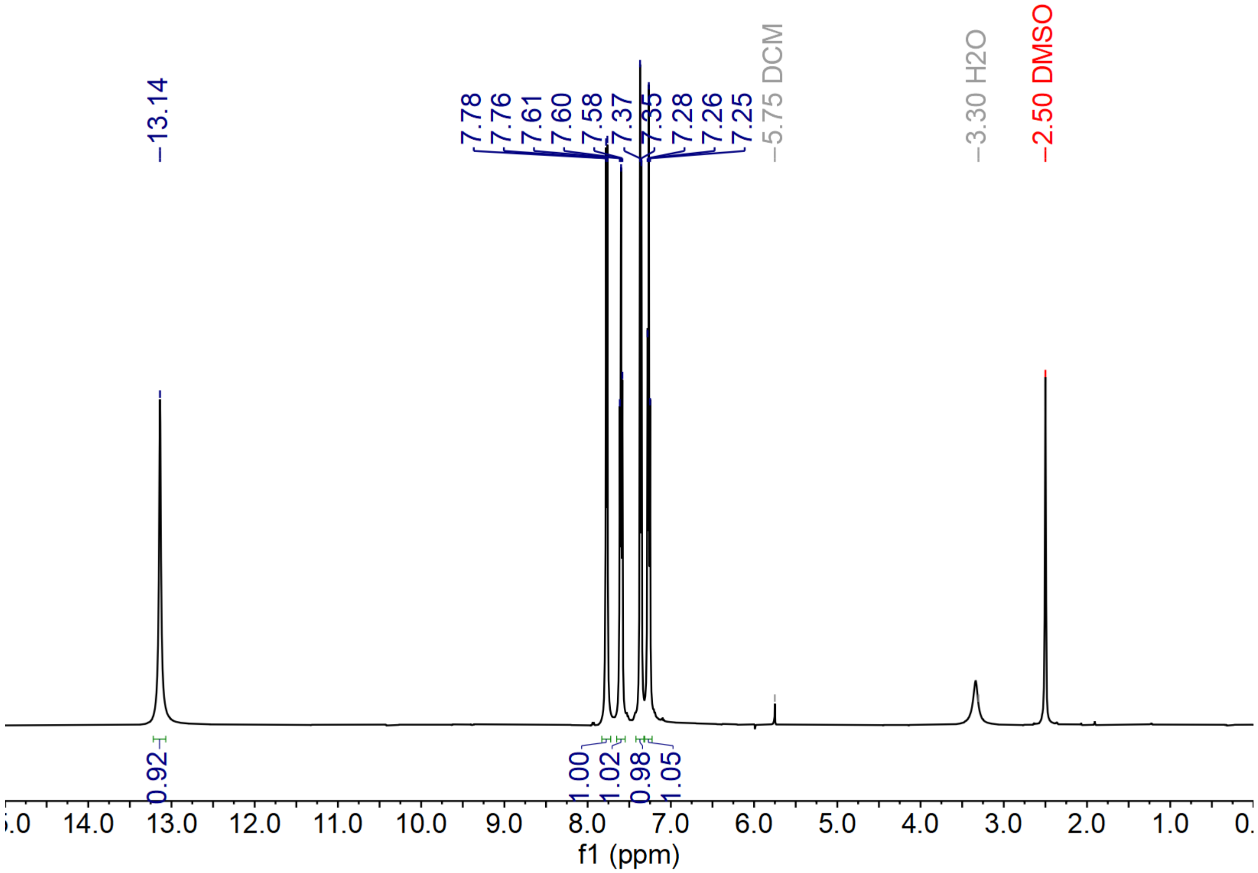


**Supplementary Figure 19.** $^{1}H$ NMR (500 MHz, DMSO-$d_6$) characterization of compound **(3)**.

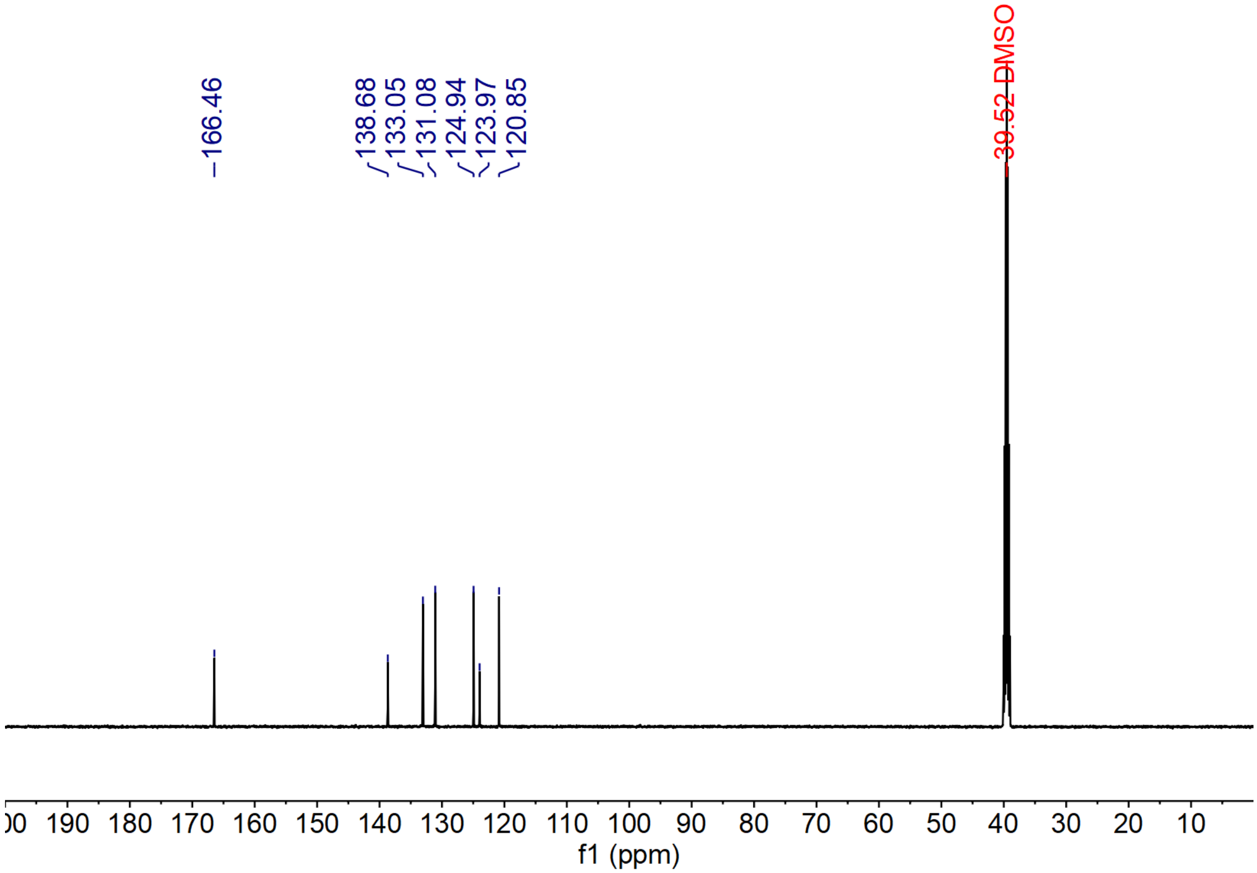


**Supplementary Figure 20.** $^{13}C$ NMR (500 MHz, DMSO-$d_6$) characterization of compound **(3)**.

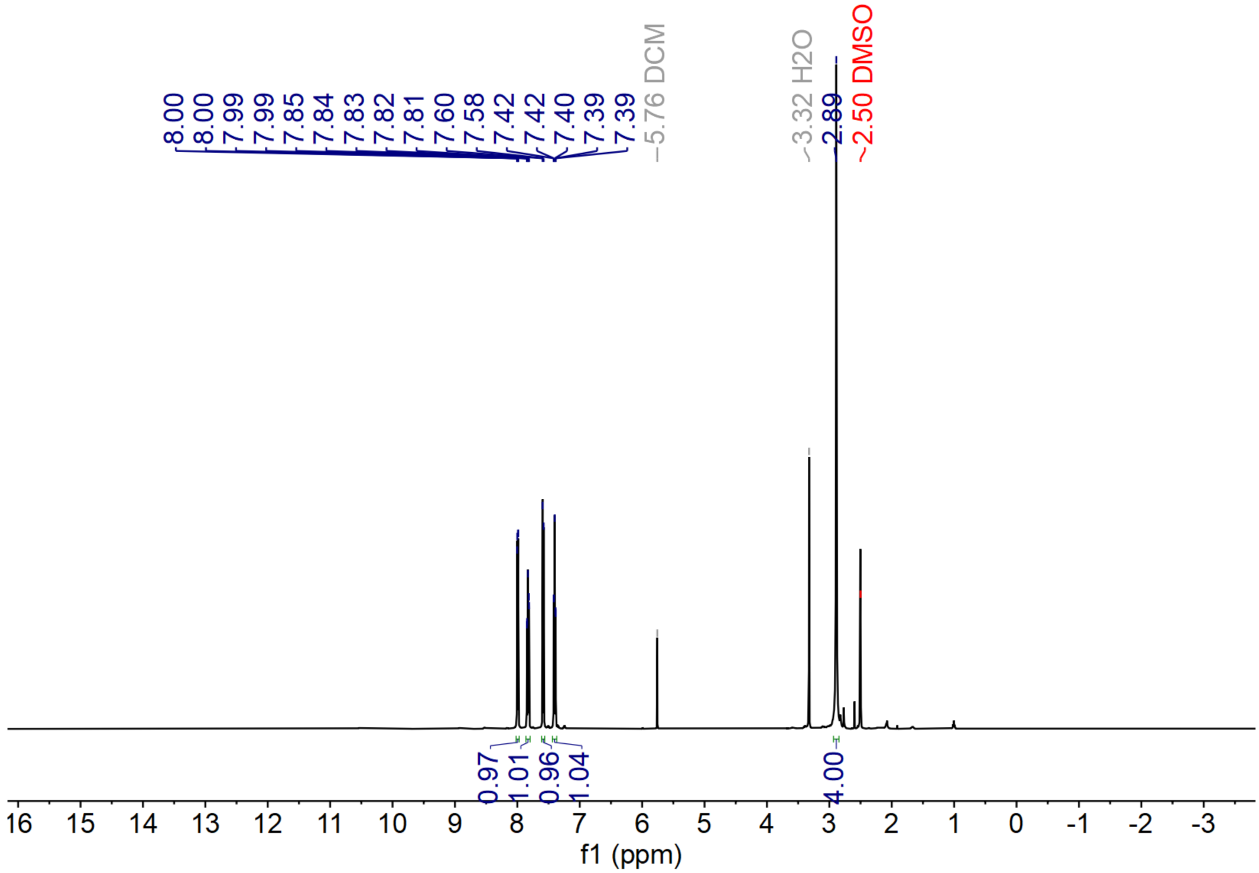


**Supplementary Figure 21.** $^{1}H$ NMR (500 MHz, DMSO-$d_6$) characterization of compound **(4)**.

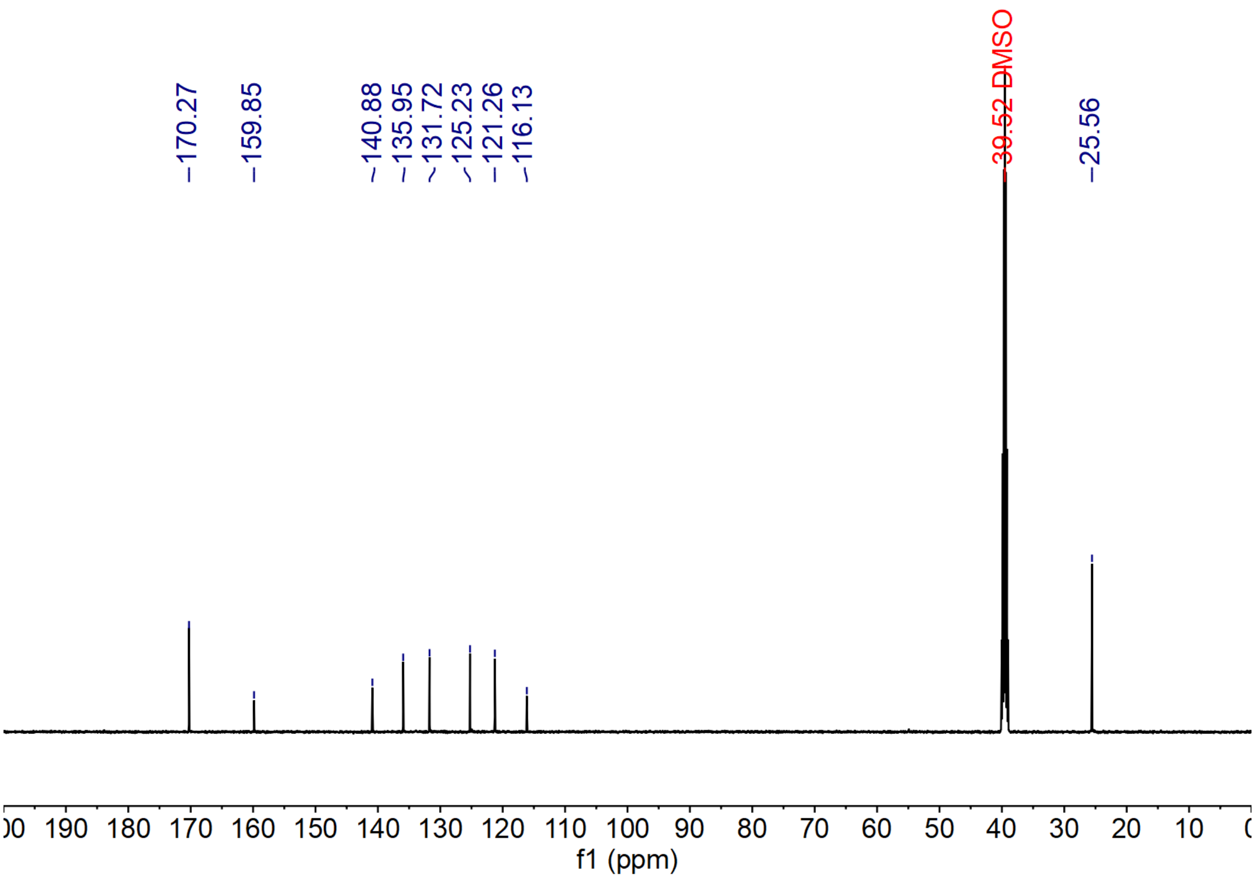


**Supplementary Figure 22.** $^{13}C$ NMR (500 MHz, DMSO-$d_6$) characterization of compound **(4)**.

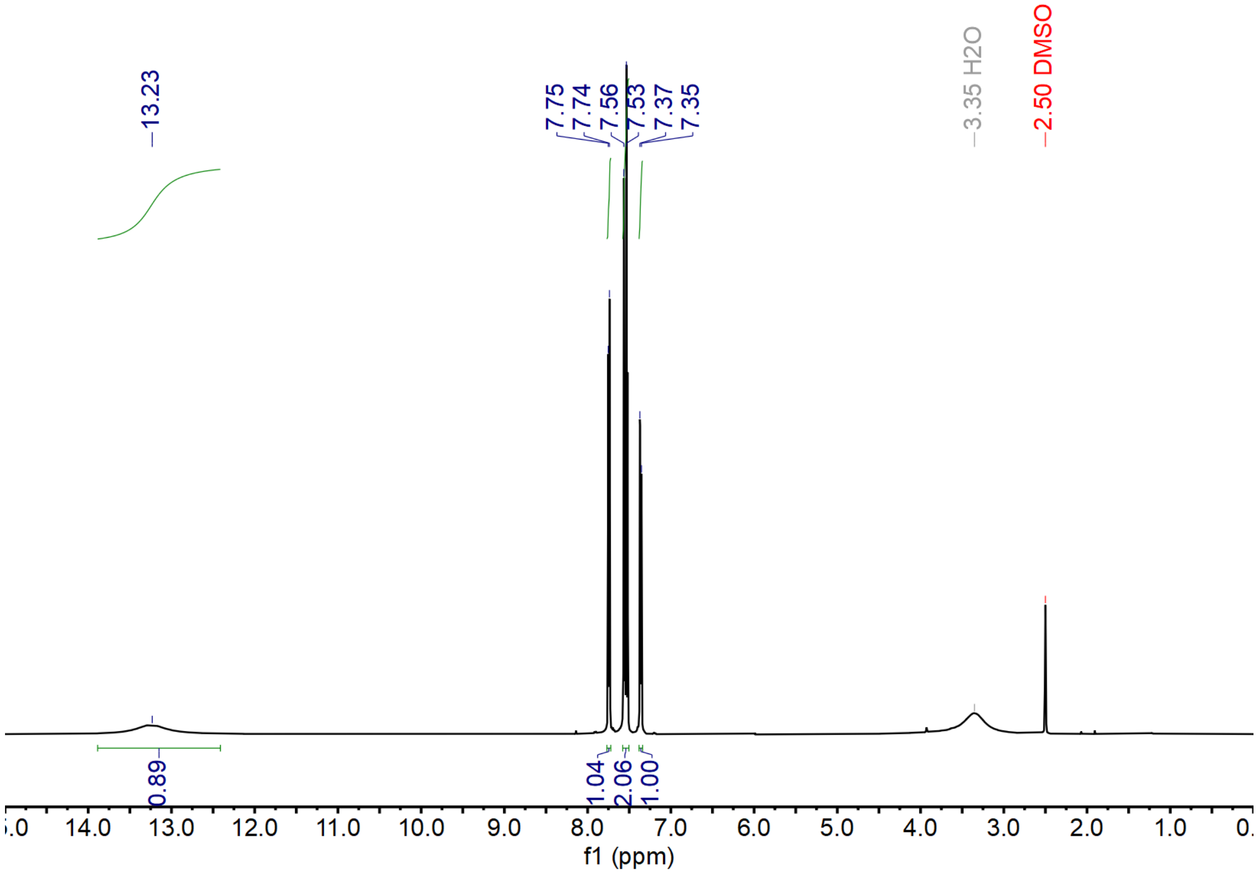


**Supplementary Figure 23.** $^{1}H$ NMR (500 MHz, DMSO-$d_6$) characterization of compound **(5)**.

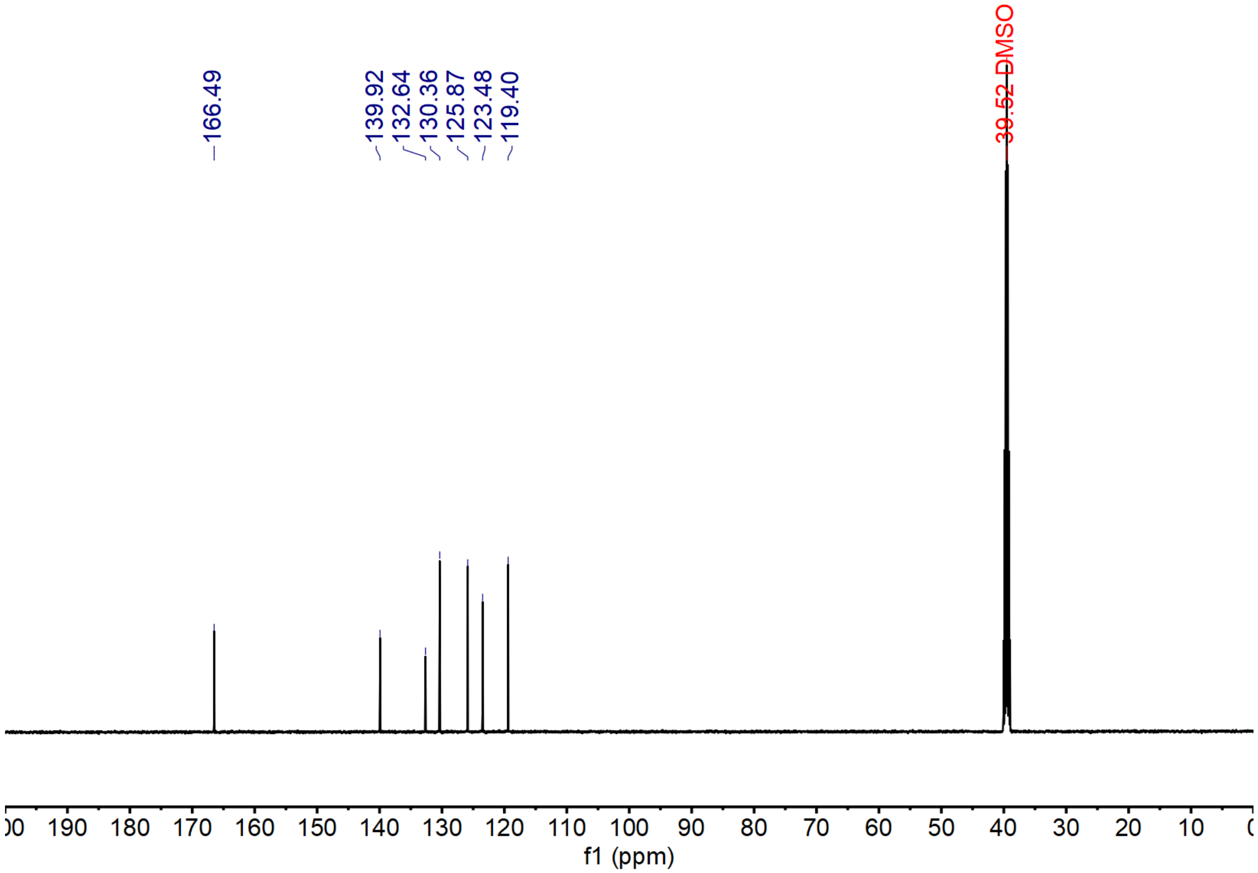


**Supplementary Figure 24.** $^{13}$C NMR (500 MHz, DMSO-$d_6$) characterization of compound **(5)**.

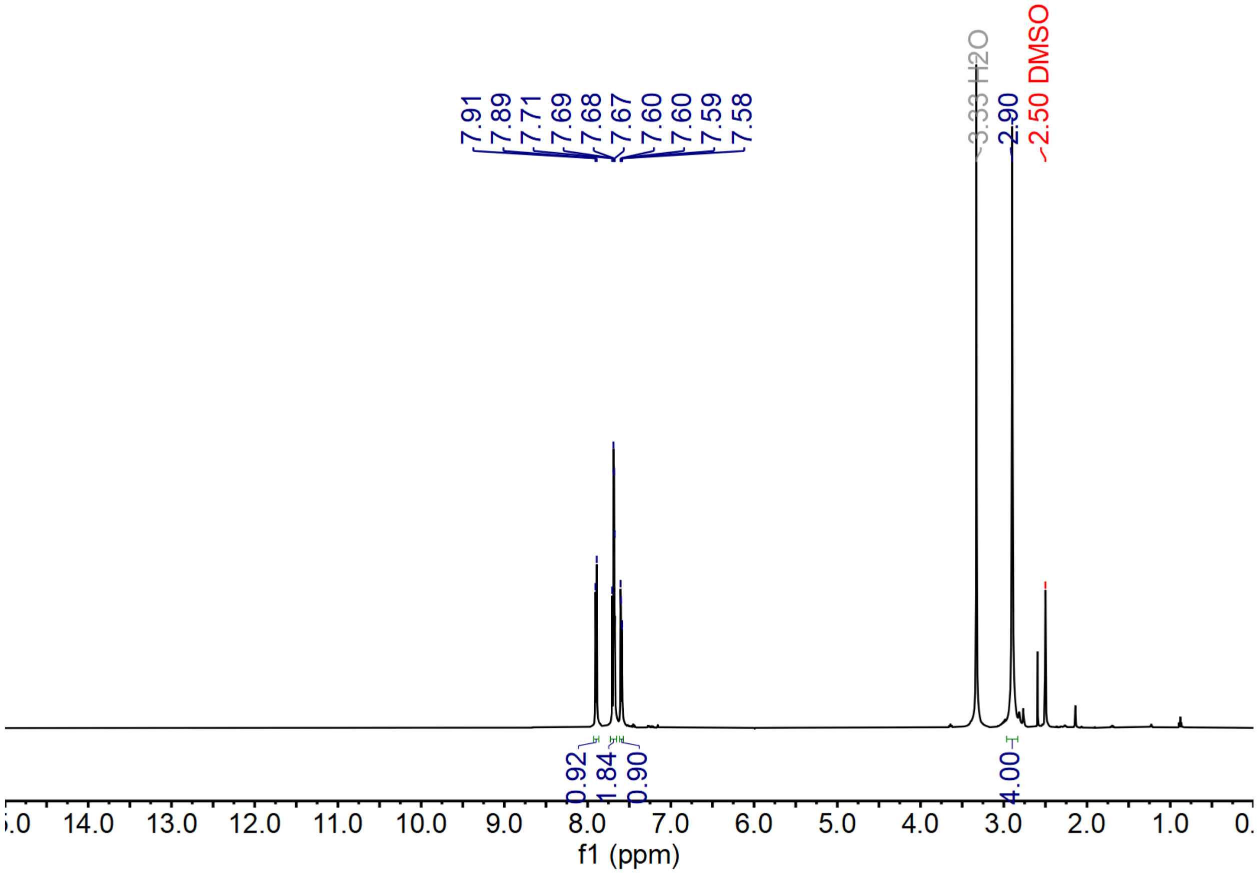


**Supplementary Figure 25.** $^{1}H$ NMR (500 MHz, DMSO-$d_6$) characterization of compound **(6)**.

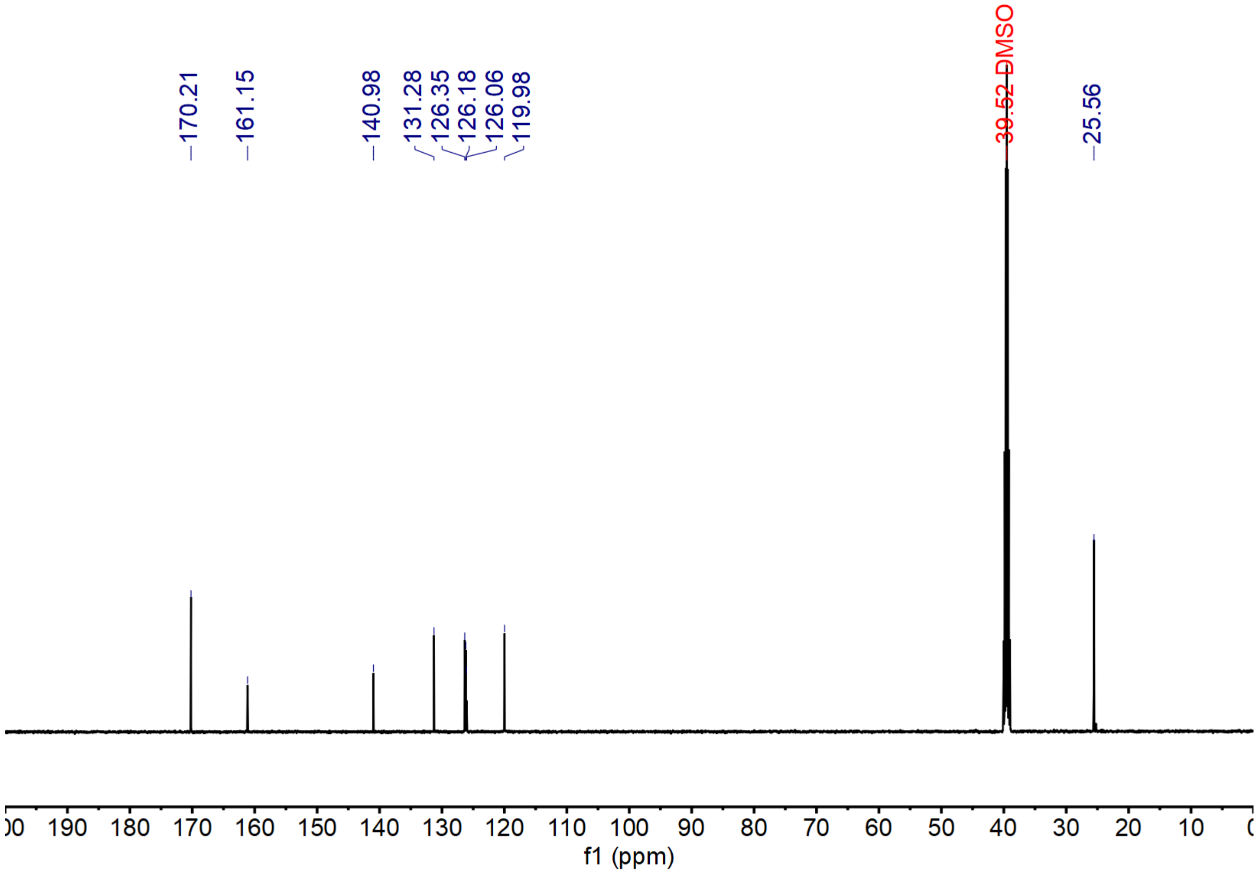


**Supplementary Figure 26.** $^{13}$C NMR (500 MHz, DMSO-$d_6$) characterization of compound **(6)**.
